# De-differentiation and Proliferation of Artery Endothelial Cells Drive Coronary Collateral Development

**DOI:** 10.1101/2022.07.27.501689

**Authors:** Gauri Arolkar, Sneha K, Hanjay Wang, Karen M. Gonzalez, Suraj Kumar, Bhavnesh Bishnoi, Pamela E. Rios Coronado, Y. Joseph Woo, Kristy Red-Horse, Soumyashree Das

## Abstract

Collateral arteries act as natural bypasses which re-route blood flow to ischemic regions and facilitate tissue regeneration. In an injured heart, neonatal artery endothelial cells orchestrate a systematic series of cellular events, which includes their outward migration, proliferation, and coalescence into fully functional collateral arteries. This process, called Artery Reassembly, aids complete cardiac regeneration in neonatal hearts, but is absent in adults. The reason for this age-dependent disparity in artery cell response is completely unknown. In this study, we investigated if regenerative potential of coronary arteries, like in cardiomyocytes, is dictated by their ability to de-differentiate. We performed single cell RNA sequencing of coronary endothelial cells and identified differences in molecular profiles of neonatal and adult endothelial cells. Neonates show significant increase in actively cycling artery cells that expressed prominent de-differentiation markers. Using both, *in silico* pathway analyses and *in vivo* experiments, we show that cell cycle re-entry of pre-existing artery cells, subsequent collateral artery formation and cardiac function is dependent on arterial VegfR2. This sub-population of de-differentiated and proliferating artery cells is absent in non-regenerative P7 or adult hearts. Together, our data indicate that adult artery endothelial cells fail to drive collateral artery development due to their limited ability to de-differentiate and proliferate.

## Background

Cardiovascular disease (CVD) represents 32% of all global deaths (World Health Organization) and its burden is increasing sharply in many new parts of the world. Earlier, deaths due to CVD dominated the western world, but currently, at least 58% of all CVD deaths occur in Asia (http://ghdx.healthdata.org/gbd-results-tool). The primary cause of CVD is coronary artery blockage, and can lead to ischemic heart disease (IHD). Despite the deadly outcomes, management of symptoms associated with IHD often include invasive procedures, which, many patients are ineligible for. Effective therapeutic approaches targeting complete cardiac regeneration, can reduce mortality and/or manage symptoms of IHD. This can be accomplished by understanding, and subsequently utilizing, the molecular mechanisms associated with different physiological aspects of IHD. One such mechanism adapted by the heart is building collateral arteries.

Large collateral arteries enhance perfusion^8^, vascularize hypoxic area, improve myocardial viability^9^, and are associated with better survival rate^10^. Collateral arteries are the arterial segments which connect a healthy arteriole to an occluded/non-perfused arteriole, and are sufficiently large to carry adequate blood flow to ischemic regions in need. Data from clinical studies suggest that the presence of native/pre-existing coronary collateral arteries present at the onset of myocardial infarction (MI) helps limit the infarct size, and improves prognosis^9,11^. The number of pre-existing collateral arteries as well as their capacity to undergo remodeling upon occlusion, are the key determinants of tissue recovery^12^. Genetic factors are proposed to regulate differences in collateral number and their functionality across human races and species^13,14^. However, a complete map of molecular mechanisms/pathways remain unknown. These molecular mechanisms could be regulated by genetic factors, and can be a useful tool to design therapeutic interventions for IHD patients.

Brain, a highly complex organ with low regenerative ability, is known to possess pre-existing collateral vessels in its pial layer. In mice, these collateral connections develop at an embryonic stage, and provide protection against ischemia in adulthood^15,16^. Pre-existing collateral arteries in mouse hind limb can grow by two distinct processes— arterialization of capillary endothelial cells (ECs)^17^ and lateral expansion/widening of smaller pre-existing collateral arteries i.e. arteriogenesis^18^. Both, in brain and hind limb, genetic background is a major contributor towards the variation in the number of observed collaterals^19,20^. Specifically, polymorphism in *Vascular Endothelial Growth Factor-a* (*Vegfa)* locus is shown to control Vegfa expression and pre-determine the extent of collateral artery network in mice^12,21^. Particularly in hind limb ischemia model, density of pre-existing collateral arteries is regulated via Vegf Receptor-1 (VegfR1)^21^. However, if VegfR1 also controls post-ischemic coronary collateral development in mice, is unknown.

In mice, complete cardiac regeneration is accomplished only when hearts are injured within a neonatal regenerative window (P0-P6). Any injury beyond this time point (>P7), limits cardiac regeneration^22,23^. Previously we have shown that, in regenerative neonatal mice, coronary collateral arteries are built by the systematic execution of 3 major cellular events driven by pre-existing artery ECs— migration, proliferation and coalescence^24^. Artery ECs expressed C-X-C motif Chemokine Receptor 4 (CXCR4), a receptor for a chemokine, C-X-C motif Chemokine Ligand 12 (Cxcl12), which steers the migration of artery ECs into the watershed. This process, named Artery Reassembly, involves disassembly of artery tips into single migratory artery cells, followed by their reassembly to build new artery segments (collateral arteries). Interestingly, despite *Cxcl12* expression in adult injured watershed, Artery Reassembly was absent in these hearts. The precise cause for absence of Artery Reassembly in adult injured hearts remains unknown.

The age-dependent discrepancy in artery EC response triggered by MI, could be because of their distinct molecular profiles^25–27^. To identify potential molecular drivers, we performed single cell RNA sequencing of neonatal and adult ECs, at a time point prior to when collateral arteries appear in the injured mouse hearts. Using both bioinformatics analyses and *in vivo* experiments, we show that, upon injury, neonatal artery ECs undergo de-differentiation and VegfR2-mediated proliferation, which is critical for collateral artery formation by Artery Reassembly and subsequent cardiac function. Both cellular events, arterial de-differentiation and proliferation, were absent in older non-regenerative hearts, post-MI. We hope, in future, these findings will aid strategizing ways to induce collateral arteries and enhance regenerative potential of cardiac tissue, by specifically targeting artery cells.

## Results

### Increase in cycling neonatal artery ECs, post-MI

Molecular profiling cardiac ECs which participate in collateral artery formation, can improve our understanding of ways in which complete cardiac reperfusion and regeneration can be attained. We thus, performed single cell RNA sequencing of neonatal post-MI heart cells. Specifically, *Cx40CreER; Rosa26^TdTomato^* mice were used, where a single dose of Tamoxifen to the lactating mothers induced expression of TdTomato fluorescence in *Cx40* expressing artery endothelial cells (aECs) in neonates. Neonates were then subjected to either sham or MI surgeries at P2, and hearts were harvested at P4 or P5 (Figure **1A**). TdTomato^+^ cardiac cells were sorted using Fluorescent Activated Cell Sorting (FACS) method, followed by sequencing using 10X genomics platform. Sequences were processed using Cell Ranger and the subsequent output files were analyzed using Seurat on RStudio^28^.

**Figure 1:**
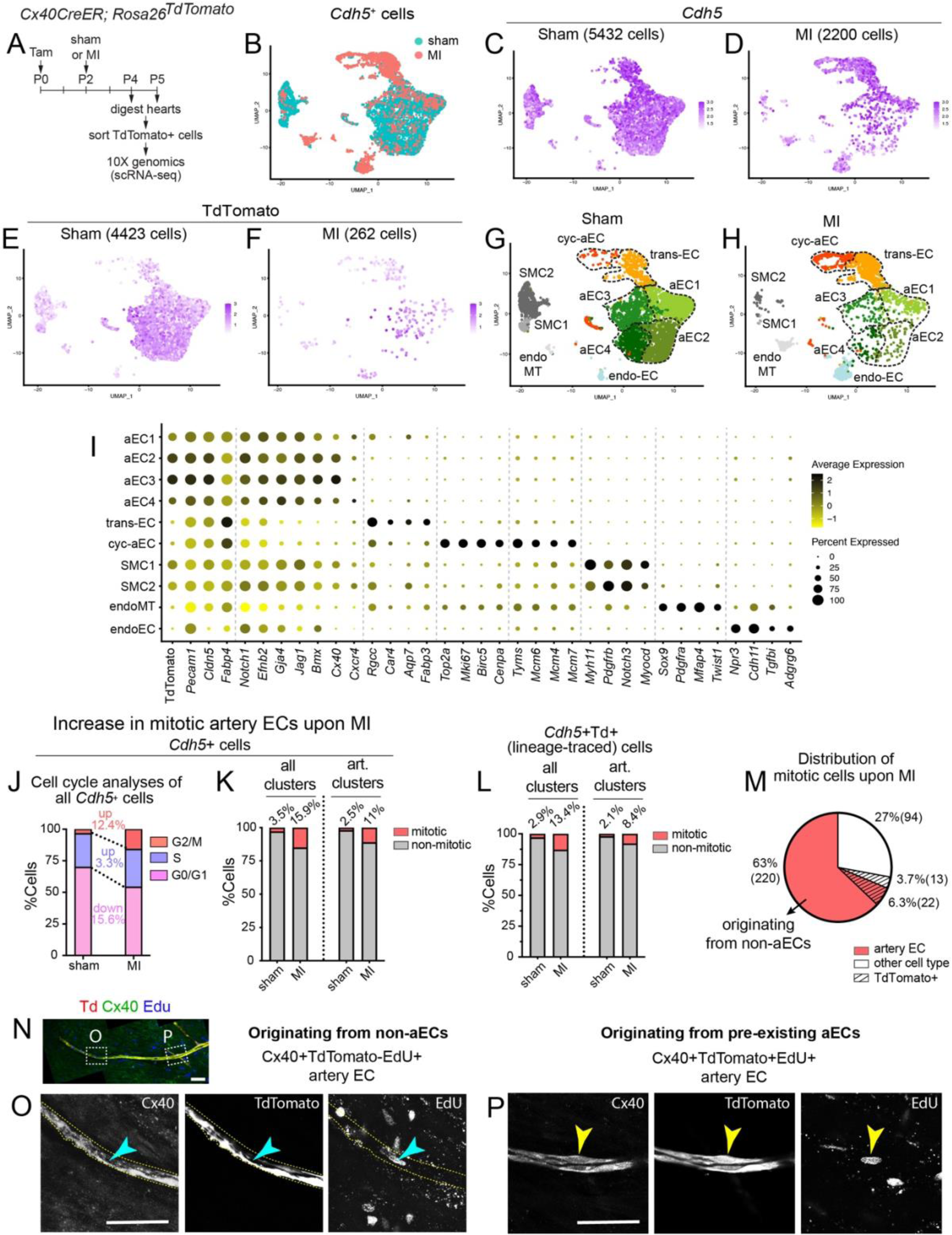
Single cell analyses of neonatal artery ECs. (**A**) Experimental setup to obtain neonatal cardiac cells for scRNA sequencing using 10X genomics (**B**) UMAP showing the distribution of *Cadherin5*^+^ neonatal sham and MI cells. (**C**, **D**) Feature Plot showing the distribution of *Cadherin5^+^* cells from (**C**) sham and (**D**) MI group. (**E**, **F**) Distribution of TdTomato expressing cells within *Cadherin5*^+^ (**E**) sham and (**F**) MI group. (**G**, **H**) UMAP showing distribution of (**G**) sham and (**H**) MI cell clusters from the merged *Cadherin5*^+^ cells. (**I**) Dot Plot showing average expression and percentage of cells, expressing relevant markers used for cluster identification. (**J**) Graph showing percentage of sham and MI *Cadherin5*^+^ cells in different cell cycle stages based on cell cycle sorting analysis on Seurat. (**K**) Graph showing percentage of *Cadherin5*^+^ cells undergoing G2 to M transition, in sham and MI groups, in all clusters or in only artery EC clusters (aEC1-4 and cyc-aEC). (**L**) Graph showing percentage of *Cadherin5*^+^ and *Cx40creER*-linegae traced (TdTomato^+^) cells undergoing G2 to M transition, in sham and MI groups, in all clusters or in only artery EC clusters (aEC1-4 and cyc-aEC). (**M**) Pie chart showing the percent distribution of G2/M *Cadherin5*^+^ cells, upon MI. (**N**-**P**) Confocal image of an (**N**) artery tip from a post-MI heart, immunostained for Cx40, lineage traced with TdTomato and labelled with EdU show presence of (**O**) Cx40^+^ TdTomato^−^ EdU^+^ (cyan arrowhead) artery EC and (**P**) Cx40^+^ TdTomato^+^ EdU^+^ (yellow arrowhead) within the same artery branch. MI, myocardial infarction; *Cdh5*, *Cadherin5;* Td, TdTomato; art, artery. Scale bar: **N**-**P**: 50μm

Sorting TdTomato^+^ cells during FACS, enriched our samples for *Cx40CreER*^+^ lineage labelled/traced artery ECs. When analyzed, these TdTomato+ cells were additionally identified to be cardiomyocytes, smooth muscle cells and cells undergoing endoMT (data not shown). We also obtained some TdTomato^−^ non-artery cells. To specifically study EC biology post-injury, we thus analyzed *Cadherin5*^+^ ECs, which included, both, TdTomato^+^ artery ECs, genetically labeled between P0-P2 (Figure **1A**), and TdTomato^−^ artery ECs (if any), identified by expression of known arterial EC genes. Pre-processing and quality control steps (Supplementary figure **S1A**) resulted in 5432 sham and 2200 MI ECs (Figure **1B**-**D**), out of which 4423 sham, and 262 MI ECs, expressed at least one copy of TdTomato (Figure **1E**, **F**). As previously observed^29,30^, compared to sham, we obtained fewer cells upon MI. This is likely due to extensive cell death following coronary ligation and subsequent tissue ischemia.

Unbiased grouping of *Cadherin5*^+^ ECs from sham and MI, resulted in 10 distinct clusters (Supplementary figure **S1B**). Merged data from sham and MI were projected onto two-dimensional space using Uniform Manifold Approximation and Projection (UMAP). The identified cell types were— mature artery ECs (aEC1-4), transitioning ECs (trans-EC), cycling artery ECs (cyc-aEC), cells undergoing endothelial to mesenchymal transition (endoMT), smooth muscle cells (SMC1,2) and endocardial ECs (endoEC) (Figure **1G, H**).

We identified the cell types with known vascular and cardiac cell markers and their relative expression levels (Figure **1I**, Supplementary figure **S1**-**3**). aEC1-4 expressed markers for mature artery ECs (*Cx40*, *Cxcr4*) and SMC1,2 expressed markers for both pericytes and SMCs (*Pdgfrb*, *Notch3*). Cells undergoing endoMT were identified by upregulated expression of mesenchymal markers such as *Col1a1* and *Cdh2* and endo-MT associated transcription factors like *Serpine1*, *Twist1*and *Snai1* (Figure **1I** and Supplementary figure **S1C**). Some of these markers were co-expressed by SMC clusters (Supplementary Figure **1C**). An EC cluster expressed several genes specific to G2/M (*Top2a*, *Mki67*, *Birc5*, *Cenpa*, *Kif2c*, *Bub1*, *Nuf2*, *Hmmr*) (Figure **1I** and Supplementary figure **S1D**-**G**) and G1/S (*Tyms*, *Mcm4*, *Mcm6*, *Mcm7*, *Ung*, *Chaf1b*, *Fen1*, *Gmnn*) (Figure **1I** and Supplementary figure **S1H-K**) transition. Sham cells in this particular cluster mostly expressed genes like *Cx40*, *Gja4*, *Jag1*, *Cxcr4* (Supplementary figure **S2A**-**D**) known to be significantly expressed and being specific to mouse^31^ and human^32^ coronary arteries. The top 20 genes in this cluster included artery specific genes^33^ like *Fgfr3*, *Slc45a4*, *Gja4* and *Ssu2* (data not shown). We hence, named the cluster as cycling aEC (cyc-aEC). EndoEC expressed *Npr3*^34^, *Cdh11*, *Tgfbi*^35^ and *Adgrg6*^36^ (Figure **1I**) and some venous markers (Supplementary figure **S1L**). Non-endothelial cardiac cells were mostly absent in our dataset (Supplementary figure **S1L**).

We also identified a cluster of cells (named as trans-EC in Figure **1G, H**) that did not express markers for any single endothelial cell sub-type. Re-clustering this group resulted into 4 distinct EC sub-clusters, those highly expressed markers for arteries, capillaries, venous/capillaries and myofibrillar genes^37^ (Supplementary figure **S3A**-**B**). This cluster especially was checked for doublets. No doublets were present in MI trans-EC cluster (data not shown). Trans-EC cluster in the sham dataset had 3 doublets (data not shown). 42.8% of artery-like ECs, in this cluster, were TdTomato^+^ (Supplementary figure **3C**) indicating their origin from pre-existing artery ECs, those labelled between P0-P2 (Figure **1A**). Additionally, this artery-like EC sub-cluster had fewer cells expressing *Cx40* or *Cxcr4* (Supplementary Figure **3B**). The indistinct molecular nature of this entire cluster, and its proximity to both, a proliferative (cyc-aEC) and mature (aEC1/3) artery EC clusters (Supplementary figure **S1B**), led us to hypothesize that cells in this cluster are in an intermediate state, i.e., between a proliferative and mature state. We hence, called this cluster, transitioning EC (trans-EC). Bioinformatics analyses of *Pecam1*^+^ cells, instead of *Cadherin5*^+^ cells, resulted in identification of the same cell types, clustered in a similar manner (Supplementary figure **S4**).

**Figure 2:**
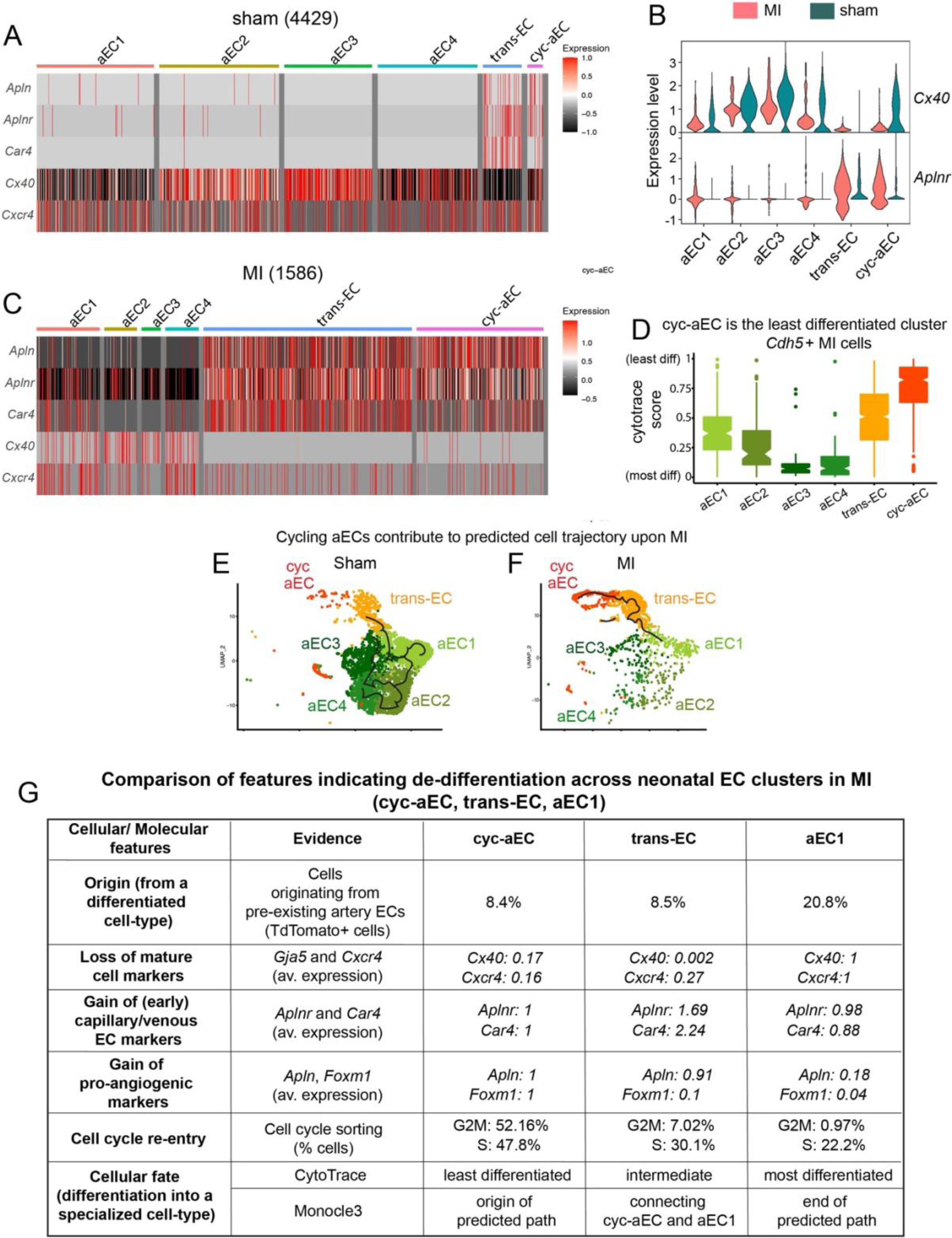
In silico assessment of de-differentiation in neonatal cycling artery ECs. (**A**) Heatmap showing gene expression of *Apln*, *Aplnr*, *Car4*, *Cx40* and *Cxcr4* across all *Cadherin5*^+^ clusters in neonatal sham group. (**B**) Violin plots comparing expression of *Cx40* and *Aplnr* between different EC clusters in sham and MI group. (**C**) Heatmap showing gene expression of *Apln*, *Aplnr*, *Car4*, *Cx40* and *Cxcr4* across all *Cadherin5*^+^ clusters in neonatal MI group. (**D**) Box plots illustrating a relative differentiation state of *Cadherin5*^+^ MI cells with scores obtained from CytoTRACE. (**E**, **F**) Trajectories obtained by performing Pseudo time analysis on neonatal *Cadherin5*^+^ (**E**) sham cells or (**F**) MI cells, using Monocle3. (**G**) Comparison of de-differentiation features across the clusters involved in pseudo time analysis of *Cadherin5*^+^ neonatal MI cells (cyc-aEC, trans-EC and aEC1).

**Figure 3:**
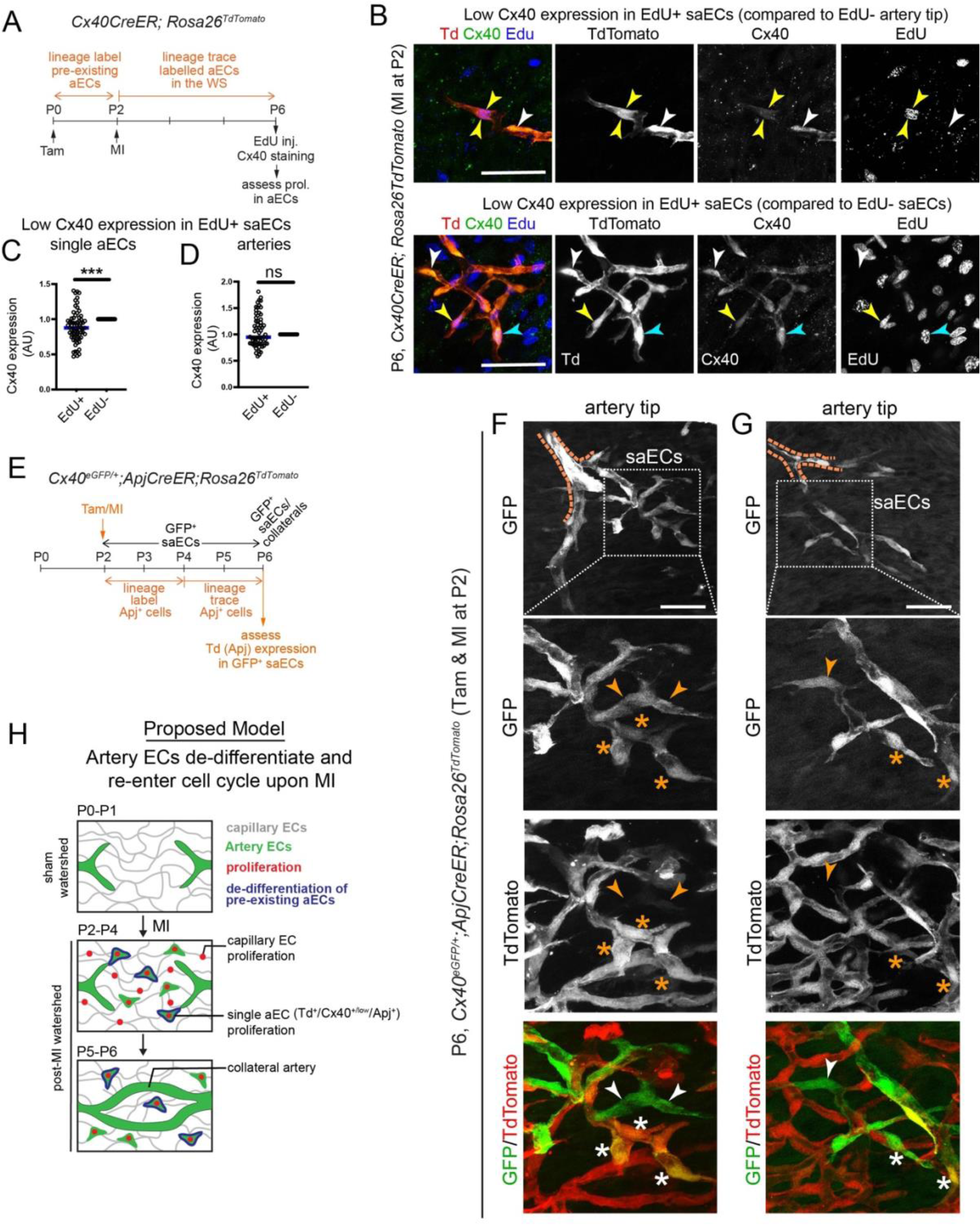
In vivo assessment of de-differentiation in neonatal cycling artery ECs. (**A**) Experimental set up to check Cx40 levels in proliferating TdTomato artery ECs post-MI (**B**) Confocal images of EdU^+^ TdTomato^+^ single artery ECs (yellow arrowhead) and EdU^−^TdTomato^+^ artery tip EC or saEC (white arrowheads) in P6 *Cx40CreER*; *Rosa26^TdTomato^* hearts, 4 days post-MI. Cyan arrowhead points to EdU^+^ Tdtomato^+^ saECs with relatively high Cx40 expression. (**C**-**D**) Quantification of cytoplasmic Cx40 in P6, *Cx40CreER*-lineage traced (**C**) single aECs (p=0.0001) and (**D**) artery ECs (p=0.09), 4 days post-MI. (**E**) Experimental set up to check presence of *Apj* in TdTomato single artery ECs post-MI. (**F**-**G**) Confocal images of GFP+ artery tips and single artery ECs showing presence (marked by *) or absence (marked by arrowhead) of TdTomato+ Apj-lineage. (**H**) A proposed model where upon MI, pre-existing stably differentiated artery ECs (green) exit artery tips as single artery ECs, de-differentiate (blue), proliferate (red) and coalesce into coronary collateral arteries. saECs are lineage traced from pre-existing artery ECs (TdTomato^+^), but downregulate expression of mature artery marker Cx40 and upregulate cap/vein EC marker Apj upon de-differentiation. Capillary ECs are shown in grey. P, postnatal day; saECs, single artery endothelial cells. Scale bar: **B**: 50μm, **F**-**G**: 50μm

Artery cells rarely proliferate, and exiting cell cycle is required for artery cell specification^24,38–40^. However, we identified an artery EC cluster (cyc-aEC) in our dataset, expressing several genes that mark G1/S and G2/M transitions (Figure **1I** **and S1D**-**K**). Cell cycle analyses of *Cadherin5^+^* cells showed a 12.4% increase in MI cells undergoing G2 to M transition as compared to the sham group (Figure **1J**). The number of MI cells in S-phase increased by 3.3% and G0/G1 phase reduced by 15.6% (Figure **1J**), suggesting a higher number of MI cells were likely mitotic.

We next performed cluster-wise quantification of mitotic cells (expressing G2/M markers), in sham and MI groups. In sham group, 2.5% of all mitotic cells were present in artery clusters (aEC1-4 and cyc-aEC) (Figure **1K**). In MI group, we observed 11% of all mitotic cells in artery clusters (Figure **1K**), indicating a 4.4-fold increase in mitotic artery ECs, upon MI. 15.9% of all *Cadherin5*^+^ cells were mitotic in the MI group, and majority of these cells (∼69.2%) were artery ECs (Figure **1K**), a stably differentiated cell type in neonatal hearts.

### MI-induces new artery development, from sources independent of pre-existing artery ECs

Post-MI artery EC proliferation is associated with Artery Reassembly and complete cardiac regeneration^24^. Use of artery EC specific *Cx40CreER* mouse line, allowed us to mark the lineage of pre-existing artery ECs with TdTomato fluorescent reporter (Figure **1A**). We performed cell cycle analysis, and quantified the number of *Cadherin5*^+^TdTomato^+^ ECs that expressed genes for G2/M transition, and were likely to be mitotic. 13.4% of all *Cadherin5*^+^TdTomato^+^ MI cells were mitotic, ∼63% of which were present in artery EC clusters (aEC1-4 and cyc-aEC) (Figure **1L**). Analyzing mitotic cell distribution within the MI group showed 69.3% of all mitotic cells to be present in artery clusters but very few (6.3%) of them were TdTomato^+^ artery ECs (Figure **1M**). This suggests that MI induces expansion of an artery EC population, not lineage labelled by, or traced from, *Cx40CreER^+^* cells. This is consistent with our previous observation that injury causes organ-wide artery growth (especially, artery-tips), independent of Artery Reassembly, and through capillary differentiation^24^.

We next performed *in vivo* experiments to confirm if an artery EC population, originating from non-artery sources (as seen in Figure **1M**) is indeed proliferating, upon MI. EdU labelled P6 hearts from *Cx40CreER; Rosa26^TdTomato^* neonates (Tamoxifen at P0, MI at P2) were harvested and immunostained for Cx40. Cx40 immunolabelled all mature artery ECs, including *Cx40CreER* lineage-traced TdTomato^+^ pre-existing aECs. We indeed, observed numerous Cx40^+^TdTomato^−^ arteries (Supplementary figure **S5A**-**C**), especially in the ischemic regions. Furthermore, we identified these artery ECs to be proliferating (EdU^+^) and integrated into Cx40^+^ vessels (Figure **1N**-**O** and Supplementary figure **S5A**-**C**). As expected, we also observed Cx40^+^TdTomato^+^EdU^+^ artery ECs (Figure **1N, P** and Supplementary figure **S5D**, **E**). Thus, ischemia can induce formation of new coronary arteries via multiple mechanisms (Supplementary figure **S5F**). Firstly, coronary collateral artery segments are generated via Artery Reassembly, a process driven by pre-existing artery ECs. Secondly, an overall organ-wide artery growth such as extension of artery tips occurs by two processes: 1) incorporation of capillary ECs as observed earlier^24^ (also suggested in Supplementary figure **S5A**-**C**) and by 2) proliferation of pre-existing (TdTomato^+^) artery ECs (as shown in Supplementary figure **S5D**, **E**).

### Neonatal artery ECs de-differentiate upon MI

Stably differentiated cells may attain a less differentiated state and re-enter cell cycle in response to pathophysiological cues. The acquired proliferative abilities allow them to transform into less specialized cell type. They can then, either re-differentiate into cells within the same lineage, or into cells of a different lineage^41^. This phenomenon can often be detected through changes in gene expression patterns. Presence of an actively cycling differentiated cell cluster in neonates (cyc-aEC) led us to hypothesize that cells in this cluster de-differentiate and contribute to new arteries, in response to MI.

De-differentiation is indicated by loss of differentiation markers, cell cycle re-entry, and expression of early genes, suggesting lack of cell fate specification. We checked if MI cells, when compared to sham cells, manifested any of these de-differentiation indicators. In sham, *Cx40*, a mature artery EC marker, was expressed by cyc-aECs, at levels comparable to mature artery clusters—aEC1-4 (Figure **2A, B**). In MI group, cells in cyc-aEC cluster expressed *Cx40*, at levels much lower than what was observed in sham cyc-aEC (Figure **2B**) or the rest of the artery ECs (aEC1-4) (Figure **2B, C**). The percentage of cells expressing *Cx40*, upon MI, reduced in all clusters (Figure **2A, C**), but this decrease was most prominent in trans-EC (19% in sham, 0% in MI) and cyc-aEC (57% in sham, 3% in MI) clusters (Supplementary figure **S6A**). A similar reduction in number of *Cxcr4*^+^ MI cells was observed specifically in cyc-aEC and trans-EC clusters (Supplementary figure **S6A**). Thus, there is an overall reduction in number of cells expressing mature artery markers, and their expression levels, in cyc-aEC and trans-EC cluster, upon MI.

During development, an immature coronary vessel plexus undergoes morphogenesis to build coronary arteries^42^. We hypothesized that cyc-aECs, if de-differentiating, would re-express some early genes expressed by this vessel plexus that mostly consists of venous/capillary ECs and from which coronaries originate during development. We hence, analyzed sham and MI cells for cardiac microvascular genes such as *Aplnr*^43^ and *Car4*^32,44^. There was a dramatic increase in number of MI cells expressing *Aplnr* in artery ECs (aEC1-4) (Figure **2A, C** and supplementary figure **S6A**), but the gene was only minimally expressed in mature artery EC clusters (aEC1-4), upon MI (Figure **2B**). In contrast to this, the total number of *Aplnr*^+^ MI cells was most prominently increased in cyc-aEC (by 2.5 fold) and trans-EC (by ∼2 fold) cluster (Supplementary figure **S6A**), and so was the expression of *Aplnr* in these clusters (Figure **2B**). *Car4* also followed the same pattern (Figure **2A, C** and Supplementary figure **S6A**), indicating both cyc-aEC and trans-EC express venous-capillary marker genes, upon injury.

Angiogenic properties of ECs facilitate vascular remodeling and subsequent cardiac regeneration. We next investigated if cyc-aEC (or trans-EC) clusters expressed any pro-angiogenic genes, yet another cellular feature of de-differentiation. Upon MI, 61% of the cyc-aECs and 47% of trans-ECs expressed pro-angiogenic *Apln* gene (Figure **2A, C** and Supplementary figure **S6A**), known to drive vascular development^45,46^ and regeneration^47–49^ *in vivo*. Expression of other angiogenic genes like *Foxm1*^50,51^, *Nrp2*^33^, *Col15a1*^33^, and *Cd34*^52^ were enriched in these two clusters (Supplementary figure **S6B**-**E**). Additionally, ECs in cyc-aEC and trans-EC clusters expressed various capillary-like/venous specific genes such as *Gpihbp1*, *Fabp3*, and *Pcdh17* (Supplementary figure **S6F**-**H**). All changes in expression of *Cxcr4*, *Cx40*, *Aplnr*, *Car4*, *Apln* in cyc-aEC between sham and MI groups, were found to be statistically significant using Wilcoxon sum rank test (**Table S1**). Thus, cyc-aEC and trans-EC clusters are likely to possess angiogenic properties, which are absent in stably differentiated artery ECs.

Cardiac vascular cells (ECs and SMCs) and cardiomyocytes have common precursor cells. In addition to classic angiogenic endothelial cell markers, cyc-aEC and trans-EC cells also expressed cardiac myofibrillar genes (*Tnnt2*, *Myl3* and *Myh6*) (Supplementary figure **S6I**-**K**). Thus, cardiac ECs maintain low, but active transcription of these cardiomyocyte specific markers as observed earlier^37^.

The differentiation status of cells and the projected pseudotime, together, can determine the root of differentiation, and hence can predict the direction of path taken by cell populations^53^. A package in RStudio ̶ Cellular (Cyto) Trajectory Reconstruction Analysis using Gene counts and Expression (CytoTRACE), uses the number of genes and their differential expression, to determine relative differentiation status of each cell with respect to others^53^. CytoTRACE score for cycling aEC, in MI dataset, was highest amongst all clusters, making it the least differentiated cluster (Figure **2D**). We also performed pseudotime analysis of sham and MI cells using monocle3 package in RStudio. In sham, the predicted trajectory connected trans-EC to aEC1-4. Sham cyc-aECs were excluded from the projected path (Figure **2E**). In contrast to sham, in MI, cyc-aEC, trans-EC and aEC1 clusters were associated with the predicted path (Figure **2F**). If multiple clusters are connected through a single trajectory, it is likely that, a less specialized cluster differentiates into a terminal cell type. This prediction is supported by our observations from genetic lineage tracing experiments *in vivo*, where upon MI, *Cx40CreER* lineage-labeled, proliferating artery ECs give rise to mature coronary collateral arteries^24^ and contribute to organ-wide artery growth (Supplementary figure **S5F**).

We next assessed the relative differentiation status of the 3 clusters participating in the trajectory obtained for MI cells (Figure **2F**)─ cyc-aEC, tans-EC and aEC1. We compared the cellular and molecular features indicating de-differentiation, across these 3 clusters (Figure **2G**). As expected, cyc-aEC and aEC1 were the least and most differentiated clusters, respectively (Figure **2G**). Trans-EC cluster showed de-differentiation features intermediate between cyc-aEC and aEC1 (Figure **2G**). Thus, it is likely that proliferating (less specialized) artery ECs in cyc-aEC cluster are differentiating into mature artery ECs in aEC1 (Figure **2F, G**). These results were supported by RNA velocity analyses, which, additionally predicts directionality of projected trajectories based on the ratio of spliced and un-spliced transcripts (Supplementary figure **S7A**, **B**). Together, we hypothesized that upon neonatal MI, at least a subpopulation of mature artery ECs originates from de-differentiation and proliferation of pre-existing/ resident artery cells.

We checked if artery ECs, those enter cell cycle upon MI, (as seen in our single cell analyses in Figure **1M**), downregulate Cx40 protein expression, in an *in vivo* setting (Figure **3A**). We compared Cx40 protein levels in EdU^+^ and EdU^−^ artery ECs labelled with Cx40 immunostaining in *Cx40CreER; Rosa26^TdTomato^* neonates, 4 days post-MI (Figure **3B**-**D**). Both categories of artery cells– single and associated with lumenized artery vessels, were analyzed. Single aECs observed in ischemic watershed are derived from resident arteries and contribute to Artery Reassembly. We observed that, Cx40 staining was significantly reduced in these EdU^+^ TdTomato^+^ proliferative single aECs (Figure **3B, C**), when compared to non-proliferative single aECs or artery tips (Figure **3B**). In contrast, EdU^+^ proliferating aECs, associated with well-formed arteries, did not show any significant change in their Cx40 expression (Figure **3D**). Together, this suggests that Cx40 is downregulated in proliferating single aECs, upon MI.

We further checked if MI-induced single artery ECs express early markers such as *Apj*, which labels capillary/venous ECs from which arteries originate during coronary development via arterialization. We used *Cx40^eGFP/+^*; *ApjCreER*; *Rosa26^TdTomato^* mice to label arteries with GFP and capillary/venous ECs with TdTomato upon Tamoxifen administration. Transgenic neonates were subjected to Tamoxifen and MI at P2, to lineage label and trace *ApjCreER*^+^ cells (Figure **3E**). Hearts were harvested at P6 when both single artery ECs and coronary collateral arteries are present (Figure **3E**) and analyzed for Apj expression in pre-existing artery ECs, upon MI. We identified single artery cells based on their location i.e. proximity to distal branches of arteries spanning the ischemic watershed area and their morphology i.e. single (not incorporated into any lumenized artery vessels) and spindle shaped as observed earlier with *Cx40CreER*-lineage trace (Figure **3B**). Indeed, many GFP^+^ single artery ECs expressed TdTomato, indicative of Apj expression at 4 days post-MI (Figure **3F** and **G**). These GFP^+^ ECs were not restricted to artery tips, but also observed in watershed regions (data not shown). Thus, several artery ECs upon MI upregulate Apj expression indicating change in cell fate in response to MI.

Together, our data suggests that single artery ECs exiting pre-existing artery tips which are the drivers of Artery Reassembly, show reduced Cx40 expression and increased Apj expression and proliferate during the course of neonatal Artery Reassembly. All three features i.e. lack of mature cell marker, increase in early angiogenic marker and cell cycle re-entry are indicative of de-differentiation. Thus, we propose that pre-existing artery ECs de-differentiate and proliferate to build coronary collateral arteries during Artery Reassembly (Figure **3H**).

### Adult artery ECs fail to proliferate during MI-induced artery neogenesis

New collateral arteries in adult mice, form in response to coronary occlusion within a week of injury^54^. However, adult coronary collateral arteries are fewer in number, have smaller diameters, and are predicted to perfuse poorly than those in neonates^55^. The reason for this age-dependent difference in which artery ECs respond to ischemia is unclear. One possibility could be that adult collateral arteries do not form naturally through Artery Reassembly. Keeping this in mind, we investigated the molecular nature of adult coronary artery ECs, and their derivative cell-types, in sham and MI mice.

Adult hearts execute Artery Reassembly only in response to exogeneous Cxcl12. As a result, artery EC lineage traced coronary collateral arteries are observed only after 14 days post-MI^24^. To capture single artery EC response that precedes Artery Reassembly, we harvested adult hearts 9 days post-MI. We performed single cell analyses of cardiac cells sorted from adult *Cx40CreER; Rosa26^TdTomato^* ventricles (Figure **4A**). Similar to neonatal data, we obtained both, TdTomato^+^ and some unlabeled cells. After performing quality control steps (Supplementary figure **S8A**), we analyzed 6256 *Cadherin5*^+^ sham and 1690 *Cadherin5*^+^ MI cells (Figure **4B, C**). Like neonatal data, our adult dataset was enriched for artery ECs (Figure **4D, E**) and we obtained fewer *Cadherin5*^+^ (Figure **3C**) and TdTomato^+^ (Figure **4F, G**) cells in MI group than sham.

**Figure 4:**
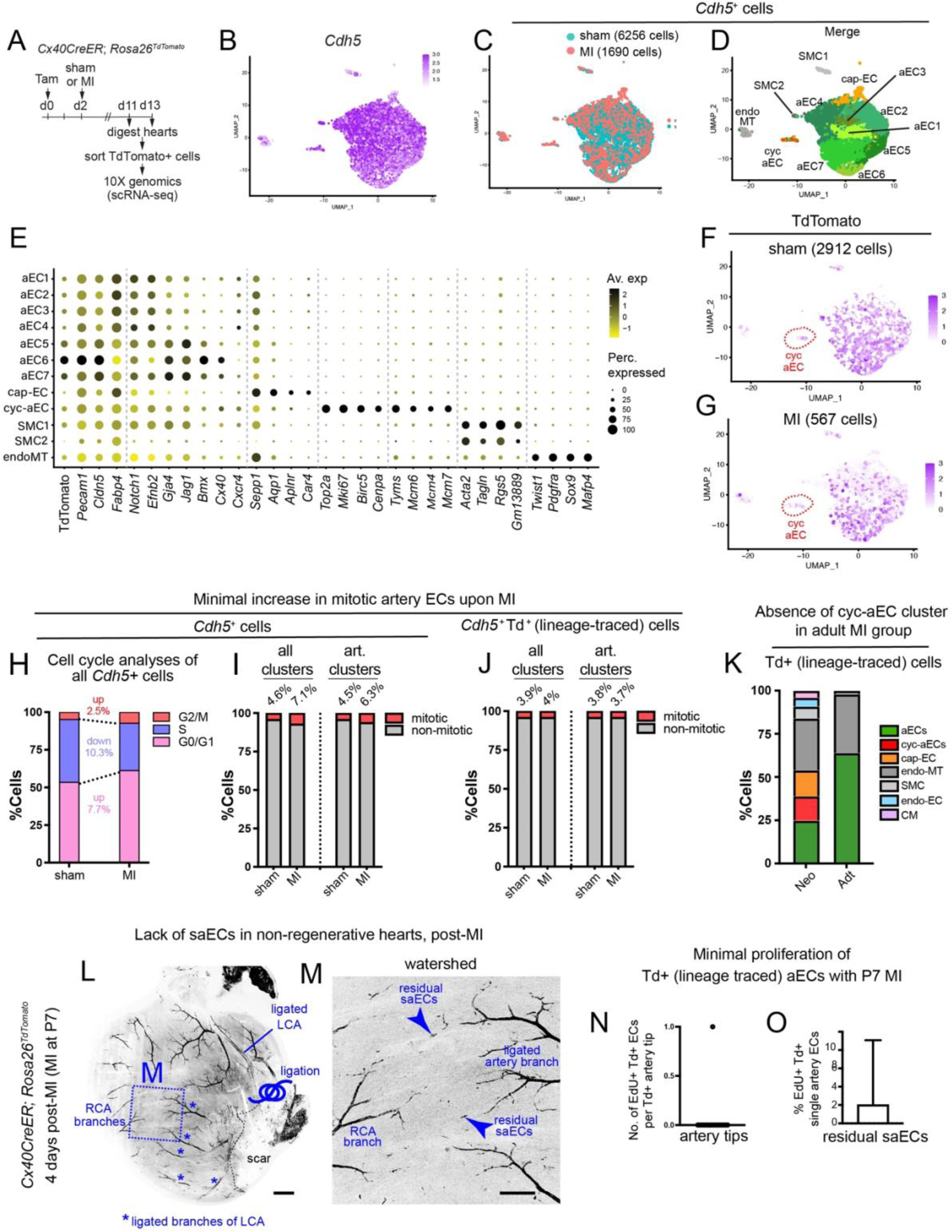
Cell cycle analyses of adult artery ECs post-MI. (**A**) Experimental setup to obtain adult cardiac cells for scRNA sequencing using 10X genomics. (**B**) Feature Plot showing the distribution of adult *Cadherin5^+^* cells from both sham and MI groups. (**C**) UMAP showing the distribution of *Cadherin5*^+^ adult sham and MI cells. (**D**) UMAP showing distribution of both sham and MI cell clusters from the merged adult *Cadherin5*^+^ cells. (**E**) DotPlot showing average expression and percentage of cells, expressing relevant markers used for cluster identification in **D**. (**F**, **G**) Distribution of TdTomato expressing cells within *Cadherin5*^+^ (**F**) sham and (**G**) MI group. (**H**) Graph showing percentage of adult, sham and MI, *Cadherin5*^+^ cells in different cell cycle stages based on cell sorting analysis on Seurat. (**I**) Graph showing percentage of *Cadherin5*^+^ cells undergoing G2 to M transition, in sham and MI groups, in all clusters or in only artery EC clusters (aEC1-7 and cyc-aEC). (**J**) Graph showing percentage of *Cadherin5*^+^ and *Cx40creER*-linegae traced (TdTomato^+^) cells undergoing G2 to M transition, in sham and MI groups, in all clusters or in only artery EC clusters (aEC1-7 and cyc-aEC). (**K**) Single cell data analyses of all TdTomato^+^ cells from neonatal and adult MI groups, showing distribution of different cell types that originate from *Cx40CreER* lineage labelling or subsequent tracing. (**L**, **M**) Representative image of P11 *Cx40CreER*; *Rosa26^TdTomato^* (**L**) whole heart and respective (**M**) watershed region, 4 days post-MI. MI was performed at P7. Lineage traced artery ECs are shown in black. * marks ligated branches of LCA, arrowheads point to “residual” saECs. (**N**, **O**) Graph showing number and percentage of EdU^+^ Tdtomato^+^ ECs in (**N**) artery tips and (**O**) residual saECs, respectively, at P11. MIs were performed at P7. P, postnatal day; LCA, left coronary artery; RCA, right coronary artery; art., artery; Adt, adult; Neo, neonate; saECs, single artery endothelial cells. Scale bars: **L**: 500μm; **M**: 200μm

Based on cell-specific gene expression patterns, we obtained 12 adult cell populations (Figure **4D**); 7 of which, constituted mature artery ECs (aEC1-7), 1 cycling artery EC (cyc-aEC), 1 capillary EC (cap-EC), 1 endoMT and 2 SMC clusters (Figure **4D, E**). Unlike neonates, trans-EC cluster was absent in adults. Instead, we identified a prominent adult capillary EC population, which when further clustered generated 3 distinct microvascular EC populations—artery-capEC, capEC, and venous-capEC (Supplementary figure **S8B**-**D**). A significant number of art-cap EC cluster (34%) expressed TdTomato fluorescence (Supplementary figure **S8D**), indicating an origin from pre-existing mature arteries. Identity of cycling aECs was confirmed with comparable expression of artery specific genes (*Gja4*, *Cx40*, *Cxcr4*, *Jag1*) in cyc-aEC and aEC1-7 (Supplementary figure **S8E**). A similar cell type distribution was also observed when bioinformatics analyses of *Pecam1*^+^ cells, instead of *Cadherin5^+^* cells, was performed (Supplementary figure **S8F**-**H**).

Adult cardiomyocytes have limited proliferation capacity^23^. To assess if the same holds true for cardiac ECs, we performed cell cycle analyses on sham and MI adult *Cadherin5*^+^ ECs. Unlike neonates, we observed only a minimal increase of 2.5% in the number of ECs undergoing G2 to M transition, upon MI (Figure **4H**). Further cluster-wise analyses revealed only 1.8% increase in the mitotic (G2/M) artery ECs (4.5% to 6.3%, Figure **4I**), compared to 8.5% increase observed in neonates (Figure **1K**), upon MI. 4.6% of adult sham cells were mitotic (Figure **4I**), suggesting that the pre-injury mitotic baseline for adult *Cadherin5*^+^ artery ECs was higher than that observed in neonates (2.5%, Figure **1K**). Thus, in comparison to neonates, fewer adult artery ECs undergo G2/M transition in response to MI.

P7 or older hearts are non-regenerative^22,23^ and do not illicit Artery Reassembly^24^. Cell cycle analyses of *Cadherin5*^+^ TdTomato^+^ lineage-traced adult ECs revealed minimal changes in G2/M ECs including in artery EC clusters (aEC1-7, cyc-aEC) (Figure **4J**). Furthermore, we checked which cell types originate from pre-existing artery ECs, upon MI. For this, we analyzed all TdTomato^+^ cells (labelled and traced from *Cx40CreER*^+^ resident artery ECs) in adult MI dataset, and compared it with neonatal MI dataset (Figure **4K**). While the neonatal MI group showed a distinct cyc-aEC cluster, this population was absent in the adult MI group (Figure **4K** and Supplementary figure **S8I**, **J**). Thus, adult hearts lack a proliferating artery EC sub-population, known to be crucial for Artery Reassembly, and complete cardiac regeneration^24^.

We next performed *in vivo* experiments to check if lack of proliferating artery cell population in injured adult hearts (Figure **4J, K**), is also observed in non-regenerative P7 hearts. *Cx40CreER; Rosa26^TdTomato^* neonates were administered with Tamoxifen to lineage label artery ECs at P5, through P7, followed by MI at P7 (Figure **4L**). Proliferation status of *Cx40CreER^+^* lineage traced, TdTomato^+^, artery ECs were analyzed with EdU label, prior to harvesting hearts at P11, 4 days post-MI. As shown earlier^24^, these hearts did not demonstrate Artery Reassembly (Figure **4L, M**), which was evident from lack of lineage traced (TdTomato^+^) collateral arteries and single artery ECs in the injured watershed regions (Figure **4M**). Very few single artery ECs were observed adjacent to artery tips (arrowheads, Figure **4M**); we called them ‘residual’ single artery ECs. EdU^+^ cells in TdTomato^+^ artery EC population—in both, artery tips (Figure **4N**) and residual single artery ECs in the watershed (Figure **4O**)—were rarely observed. Together, data from both bioinformatics analyses and *in vivo* experiments indicate that artery ECs in non-regenerative P7 or adult hearts have limited self-renewing capacities.

### Adult artery ECs do not de-differentiate, upon MI

We next checked if adult MI *Cadherin5*^+^ cycling aECs (Figure **3I**) demonstrate any characteristics indicative of de-differentiation. Though adult MI cycling aEC cluster was the least differentiated aEC cluster (Figure **5A, B**), its CytoTRACE score was comparable to a mature artery cluster—aEC5 in MI (Figure **5B**). Expression of proangiogenic marker *Apln*, and its receptor *Aplnr* were seemingly upregulated in this cluster, but there was no change in expression of *Car4*, *Cx40*, or *Cxcr4* (Figure **5C**). We analyzed the cellular and molecular features of neonatal cyc-aECs, which were indicative of de-differentiation, and checked those indicators in adult cyc-aECs (Figure **5D**). Some of the key de-differentiation features absent in adult cyc-aECs were—1. their adequate number, which is representative of their ability to self-renew/ re-enter cell cycle 2. loss of mature cell markers and 3. gain of microvascular/ pro-angiogenic markers (shown in red, Figure **5D**). For instance, in contrast to neonatal MI cyc-aECs, adult MI cyc-aECs, showed almost no change in the number of *Cx40* expressing cells (Figure **5D**). Adult cells expressing *Cxcr4* in this group, increased by 1.76 fold (Figure **5D**). The number of cells expressing *Apln*, *Aplnr*, and *Car4*in the adult cyc-aECs were increased upon MI. However, these changes in cell numbers or gene expression levels, were statistically insignificant when compared between adult sham and MI groups, using Wilcoxon sum rank test (* in Figure **5D** and **Table S1**). Furthermore, cycling aEC cluster did not contribute to the predicted path associated with mature aECs (aEC1-7), even in MI group (Figure **5E, F**). Thus, unlike neonate, adult cycling aECs were neither connected to artery ECs in trajectory analyses (Figure **5E, F**), nor were lineage traced from pre-existing arteries (Figure **4J, K**). Together, adult cycling aEC population fails to drive artery regeneration (Figure **5G**) due to its limited number and inability to de-differentiate (Figure **5D**).

**Figure 5:**
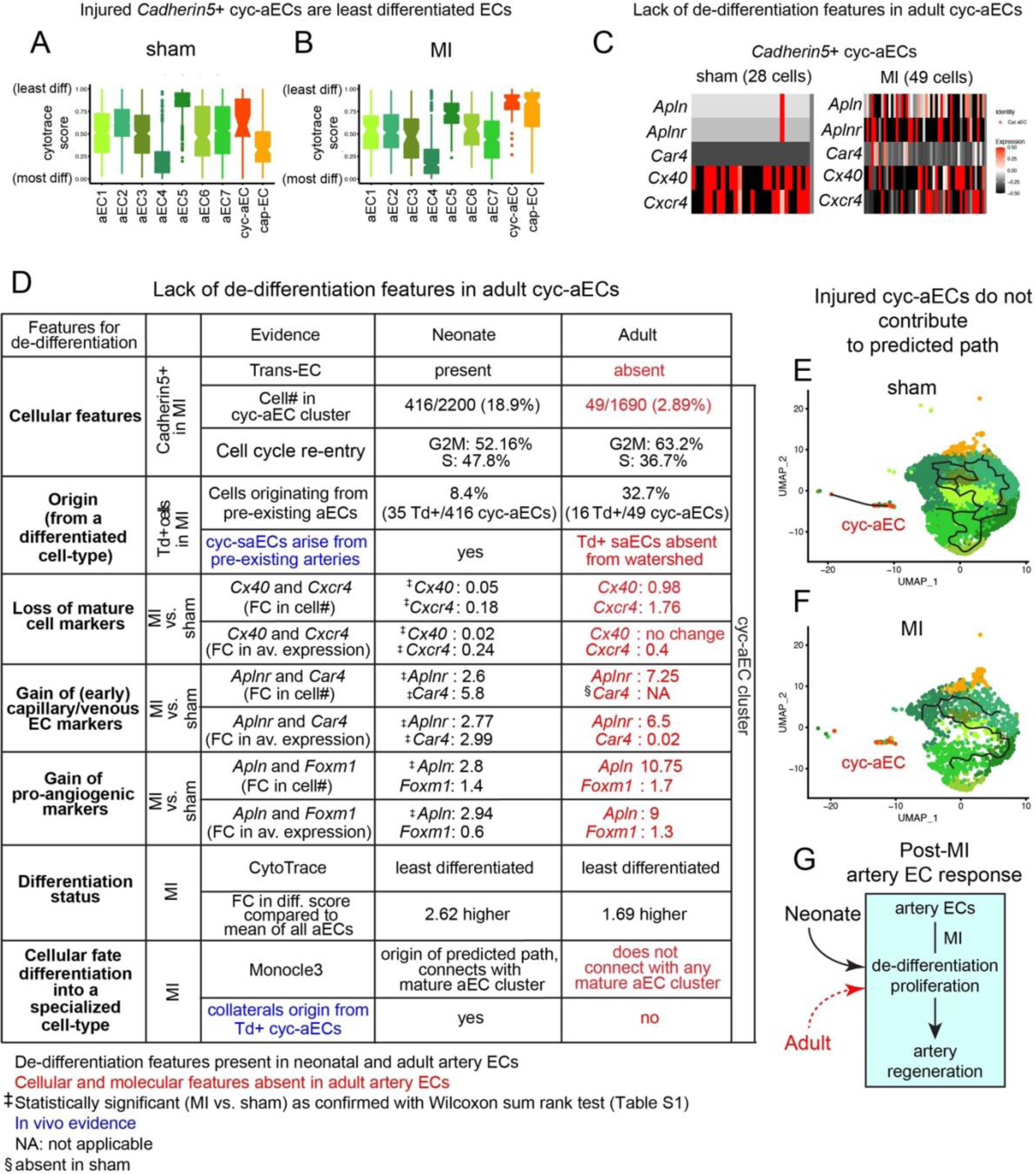
Analyses of de-differentiation features in adult MI cells. (**A**-**B**) Box plots illustrating a relative differentiation state of adult *Cadherin5*^+^ (**A**) sham and (**B**) MI cells, with scores obtained from CytoTRACE. (**C**) Heatmap of adult sham and MI cells showing the expression of *Apln*, *Aplnr*, *Car4*, *Cx40* and *Cxcr4*, in their respective cyc-aEC cluster. (**D**) Comparison of de-differentiation features in *Cadherin5*^+^ neonatal and adult cyc-aEC cluster. (**E**-**F**) Pseudo time analysis of adult *Cadherin5*^+^ cells from (**E**) sham and (**F**) MI group. (**G**) Schematic showing post-MI neonatal artery ECs accomplish artery regeneration by undergoing de-differentiation and proliferation; a phenomenon rarely observed in adult. Dotted line, minimal contribution. FC, Fold Change; av., average; diff, differentiation; Td, TdTomato; #, number.

### Arterial VegfR2 promotes artery EC proliferation during Artery Reassembly

Pre-existing artery ECs drive collateral artery formation in neonatal mice, only upon MI^24,56^. We isolated all genes from *Cx40CreER*-lineage traced TdTomato^+^ injured artery ECs (Figure **1F**) and performed gene ontology enrichment analysis and visualization (GOrilla). This allowed us to identify biological processes triggered upon MI. Key vascular responses such as angiogenesis, migration and proliferation, were significantly upregulated upon MI (Figure **6A**). Artery Reassembly is a systematic step-wise cellular event where artery EC migration and proliferation are crucial. Thus, our neonatal single cell dataset contains the artery EC population, including those that drive Artery Reassembly, and subsequent neonatal cardiac regeneration.

**Figure 6:**
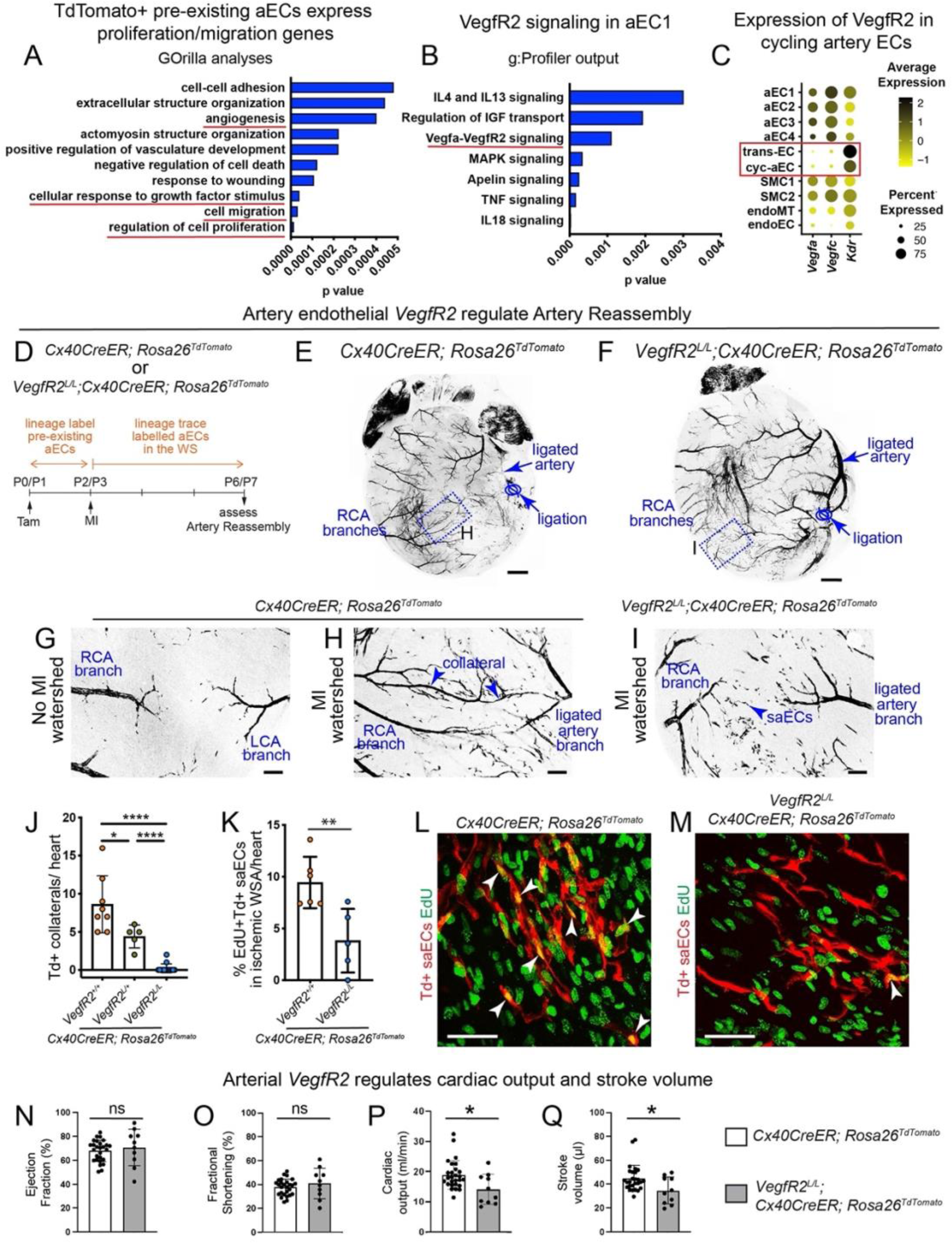
Vegf pathway in neonatal Artery Reassembly and cardiac function. (**A**) Results from Gene Ontology analysis on TdTomato^+^ cells from neonatal MI group (with sham as background), using the tool GOrilla. (**B**) Pathway analysis on genes expressed in neonatal aEC1 cluster, using g:Profiler. (**C**) DotPlot showing average expression and percentage of cells expressing *Vegfa*, *Vegfc* and *Kdr* (*VegfR2*) in all *Cadherin5*^+^ cells. (**D**) Experimental set up to test the role of arterial VegfR2 in neonatal Artery Reassembly. (**E**, **F**) Representative image of (**E**) control and (**F**) *VegfR2* artery endothelial cell specific knockout whole hearts, 4 days post-MI. (**G**-**I**) Representative confocal images of P6 watershed regions from (**G**) un-injured, (**H**) MI control and (I) MI *VegfR2* knockout hearts. MIs were performed at P2. Lineage traced artery ECs are shown in black. (**J**) Quantification of *Cx40CreER*-lineage traced collateral artery numbers observed 4 days post-MI. * represents p-value 0.0368, **** represents p-value <0.0001 (**K**) Quantification of percentage of EdU^+^ Tdtomato^+^ saECs in watershed regions from *Cx40CreER*-lineage traced hearts, 4 days post-MI. ** represents p-value 0.0083 (**L**, **M**) Representative confocal images of EdU^+^ Tdtomato^+^ saECs (white arrowheads) in watershed regions from *Cx40CreER*-lineage traced hearts, (**L**) with or (**M**) without arterial *VegfR2*, 4 days post-MI. LCA, left coronary artery; RCA, right coronary artery; WSA, watershed Area. (**N**-**Q**) Graphs showing various parameters of cardiac function as measured in adult wild-type (n=27) and *VegfR2* artery-EC specific knockout mice (n=10). p=0.5067 (Ejection Fraction), p=0.3401 (Fractional Shortening), p=0.0108 (Cardiac Output), p=0.0186 (Stroke Volume). Scale bars: **E**, **F**: 500µm; **G**-**I:** 100 µm; **L**, **M**: 50 µm

aEC1, in neonatal MI dataset, is the only artery EC cluster associated with the predicted trajectory and connected with cycling artery ECs (Figure **2F**). We performed pathway analyses on *Cadherin5*^+^ aEC1 cells using g:Profiler and filtered pathways from various databases, based on their statistical significance (p>0.05). Many pathways relevant to vascular development and injury response were enriched in aEC1 such as Apelin, TNF and Interleukin signaling^57^ (Figure **6B**). Vegfa-VegfR2 signaling (p=0.0011) was of particular interest (Figure **6B**), as it has established molecular functions in mouse coronary artery development^40,58^ and neovascularization upon injury in different tissues^59,60^.

We checked the expression levels of ligands (*Vegfa*, *Vegfc*) and a major receptor (*VegfR2* or *Kdr*) associated with coronary morphogenesis, in our dataset. While *Vegfa* and *Vegfc* were expressed in artery ECs (aEC1-4), *VegfR2* was specifically enriched in cycling aEC and trans-EC population (Figure **6C**). The *VegfR2* expression, was significantly different in cyc-aECs, when compared between neonatal sham and MI groups, using Wilcoxon sum rank test (**Table S1**). When immunostained for VegfR2, the protein was located in coronary artery ECs of small arteries, artery tips and capillary ECs (Supplementary figure **S9A**). The levels of VegfR2 protein in arteries was lower as compared to capillaries, and the protein was almost always absent in large arteries (Supplementary figure **S9A**). Both EC types (arteries and capillaries) facilitate Artery Reassembly, either by directly participating in the process (artery ECs) or providing the Cxcl12 ligand (capillary ECs). Given VegfR2 expression in both, capillaries and small arteries, we hypothesized that Vegf pathway has a role in Artery Reassembly, post-MI.

We analyzed the role of arterial *VegfR2* in Artery Reassembly, *in vivo*. Control (*Cx40CreER*; *Rosa26^TdTomato^*) and artery EC-specific, conditional *VegfR2* knockout (*VegfR2^L/L^*; *Cx40CreER*; *Rosa26^TdTomato^*) neonates received Tamoxifen at P0/P1 which allowed labelling of pre-existing artery ECs. These hearts were subjected to MI at P2/P3 and hearts were harvested 4 days post-MI, at P6/P7 (Figure **6D**). Whole heart imaging allowed us to visualize the entire arterial network of the heart (Figure **6E, F**) as well as focus into regions of interest such as the watershed region where new collaterals are expected to appear after injury (Figure **6G**-**I**). Control and *VegfR2* artery specific knockout hearts did not illustrate any gross differences in their baseline artery architecture, without MI (Supplementary figure **S9B**-**D**). Watershed regions from sham (data not shown) or no injury (Figure **6G**) hearts, did not show any collateral arteries lineage labelled with TdTomato, suggesting absence of Artery Reassembly in non-ischemic hearts. Upon MI, control hearts consistently showed several (average of 8) *Cx40CreER*-lineage traced (TdTomato^+^) collateral arteries as observed earlier^24^ (Figure **6E, H**). In contrast, we rarely observed any such collateral arteries in *VegfR2* artery specific knockout hearts (Figure **6F, I, J**). This suggests an artery specific role of VegfR2 in executing Artery Reassembly, upon MI.

We next investigated the cause for lack of TdTomato^+^ collateral arteries in *VegfR2* knockout watershed regions. Artery Reassembly is a three step cellular event—migration, proliferation and coalescence, of artery ECs. We first assessed the ability of TdTomato^+^ cells to migrate into the watershed. Qualitative and quantitative observations suggested that the number of *Cx40CreER*-lineage traced single artery ECs, observed in the watershed regions, were comparable in control and knockout MI hearts (Figure **6H, I** and Supplementary figure **S9E**-**H**). This suggests that in absence of arterial VegfR2, the migration of artery ECs from their resident arteries into the watershed region is less likely affected. We next investigated the proliferation step, and injected neonates with EdU, one hour prior to harvesting the hearts. We then, quantified the number of proliferating single aECs in the MI watershed regions. Interestingly, *VegfR2* knockout watershed regions showed a ∼2.4-fold reduction in the number of proliferating single artery ECs as compared to control hearts (Figure **6K**-**M**). Such reduction in number of proliferating single artery ECs was also observed in non-regenerative P7 injured hearts (Figure **4O**). Thus, arterial VegfR2 regulates neonatal artery EC proliferation, post-MI. A defect in VegfR2-mediated artery cell proliferation could also affect subsequent coalescence of TdTomato^+^ cells and successful execution of Artery Reassembly, as observed in *VegfR2* knockout hearts (Figure **6J**).

Functional coronary collateral arteries built by Artery Reassembly promote cardiac regeneration. Since arterial deletion of *VegfR2* reduced number of collateral arteries, we tested if this neonatal *VegfR2* deletion affects cardiac function in adults. We performed MIs a neonatal stage (P2) and checked several parameters for cardiac function 30 days post-MI. While parameters like Ejection Fraction and Fractional Shortening did not change significantly (Figure **6N, O**), cardiac output and stroke volume were significantly reduced (Figure **6P, Q**) in *VegfR2* arterial EC specific knockouts. Thus, arterial VegfR2 facilitates neonatal Artery Reassembly which regulates cardiac function in adult mice.

We next investigated if Vegf ligands and receptors are expressed in adult ECs. The expression of Vegf ligands, *Vegfa* and *Vegfc*, were mostly observed in mature artery EC clusters (Supplementary figure **S10A**) like in neonatal data (Figure **6C**). The receptor *VegfR2* (*Kdr*), was upregulated in capillary ECs, as expected (Supplementary figure **S10A**). Interestingly, at least at the mRNA level, *VegfR2* was downregulated in ∼75% of actively cycling aECs (Supplementary figure **S10A**). This is in contrast to the upregulated expression of *VegfR2* in neonatal cyc-aECs (Figure **6C**). When compared to sham group, *VegfR2* was significantly downregulated in adult MI cyc-aEC cluster (**Table S1**). Given the functional significance of arterial *VegfR2* in neonatal aEC proliferation (Figure **6K**-**M**), the lack of *VegfR2* expression in “residual” adult cycling aECs, could explain depleted potential for aEC proliferation in injured adults. In summary, our data indicate that arterial VegfR2 drives, specifically, the proliferation step of Artery Reassembly in injured neonatal mice (Supplementary figure **S10B**).

## Discussion

In this study we utilize both *in silico* analyses and *in vivo* experiments to explain the age-dependent artery EC responses to cardiac injury. Firstly, with single cell sequencing data analyses we show, for the first time, that coronary artery ECs, an otherwise stably differentiated cardiac cell type, undergoes injury-induced de-differentiation in neonates, but not in adults. In this study, we defined artery EC de-differentiation as a biological process involving four major cellular/molecular features 1. downregulation mature artery EC cell markers, 2. expression of early and angiogenic markers, 3. cell cycle re-entry and 4. Subsequent differentiation into more artery ECs to build new collateral arteries by Artery Reassembly. With the combination of bioinformatics and mouse experiments, we show that the ability of artery ECs to de-differentiate, correlates with their ability to proliferate. Both cellular responses ↓artery cell de-differentiation and proliferation↓ are specific to neonatal regenerative window. Secondly, using *in vivo* experiments, we discover a second artery-specific receptor (apart from Cxcr4, which controls artery cell migration during Artery Reassembly^24^), which regulates proliferation of artery ECs in response to neonatal MI. Expression of this receptor, *VegfR2*, is downregulated in adult cycling artery ECs and could explain absence of Artery Reassembly in adults. Finally, our data suggests that each step of Artery Reassembly is regulated by a distinct signaling axis, and the successful execution of each step will require activation of these molecular pathways in the right space and time. In summary, we highlight the differences in age-dependent molecular profiles of mouse coronaries and propose a mechanism for reduced plasticity of artery ECs and lack of Artery Reassembly in non-regenerative P7 or adult hearts.

We used single cell RNA sequencing to investigate MI-induced artery EC response in neonates and adults. We obtained fewer artery ECs upon MI as compared to sham. This was observed in both neonates and adults. In our dataset, the time points when the neonatal or adult hearts were harvested represent a state when cardiac regeneration has not been completed yet. At the chosen timepoints, while many vascular regenerative processes have been initiated, it is likely that cell death would still be in progression. This phenomenon is also reflected in *Cx40CreER*-lineage traced TdTomato artery ECs. Such small fraction of artery ECs observed in our dataset have also been reported in other neonatal and adult^27^ datasets^29,30^ and likely represent extensive cell death following coronary ligation and a vascular environment prior to complete cardiac recovery.

TdTomato cells in our MI datasets were ∼17 fold less in neonates and ∼5 fold less in adults as compared to their sham counterparts. We hence analyzed all ECs which included both TdTomato+ and TdTomato-cells. The analyses of *Cadherin5*+ all ECs allowed us to build several hypotheses which were tested in vivo. Using genetic lineage tracing and confocal imaging of whole hearts, we show that post-MI, 1) neonatal artery ECs also originate from sources other than pre-existing artery ECs. 2) During Artery Reassembly cycling pre-existing artery ECs downregulate mature artery EC marker (*Cx40*) and upregulate marker for capillary/vein (*Apj*) from which artery ECs originate during development 3) artery EC proliferation is minimal in non-regenerative P7 hearts when subjected to MI and finally, 4) injury induced artery EC proliferation is regulated by VegfR2 mediated pathway which subsequently impacts cardiac function of adult mice. These in vivo results were based on the findings from our single cell dataset. Thus, despite of their low numbers in our single cell datasets, the artery EC population is representative of post-injury vascular niche in vivo.

### De-differentiation and Proliferation are Coupled during Cardiac Regeneration

New cardiomyocytes originate from pre-existing cardiomyocytes upon injury^22,23^ by de-differentiating and re-entering cell cycle^61,62^. Cardiomyocyte de-differentiation has been elucidated upon adult zebrafish cardiac injury via disassembly of sarcomeric structures, detachment of cardiomyocytes from one another, and expression of early embryonic genes (α-Smooth Muscle Actin (αSMA), embryonic Myosin Heavy Chain, and Gata4)^61–63^. Post-MI adult cardiomyocytes also demonstrate somewhat limited de-differentiation potential indicated by downregulation of mature cardiomyocyte functions such as hypertrophy, contractility, electrical conduction, and upregulation of several early genes like *αSMA*, *Gata4*, *Runx1* and *Dab2*^64^. Unlike cardiomyocytes, EC de-differentiation has not been reported in any cardiac injury model. We show here, for the first time, that neonatal mouse coronary artery ECs undergo de-differentiation upon MI; a process limited in adults.

Since there are no objective indicators of artery EC pluripotency, one can argue that changes in gene expression or phenotypes could be a reflection of their plasticity, and not necessarily reversion of cell fate into a de-differentiated state. It is also possible that number of de-differentiated cells observed in cyc-aEC and transEC clusters of uninjured hearts simply increase upon MI. To address this, we combine single cell analyses and genetic lineage tracing. Lineage labelled non-cycling artery ECs in MI hearts can be sequentially traced into cycling single artery ECs in the watershed, and consequently into non-cycling artery ECs on the coronary collaterals. We analyzed the transiently cycling artery ECs and show that several indicators for de-differentiation, were enhanced in neonates. One such indicator was presence of Apj lineage in Cx40^+^ artery ECs. It is possible that Apj-lineage traced capillaries differentiate into artery ECs, a phenomenon called arterialization already shown to contribute (though minimally) to collateral formation by Artery Reassembly^24^. Our in vivo data cannot distinguish between differentiation of capillary ECs into artery ECs and upregulation of capillary marker (Apj) in arteries. However, the unique location and morphology of cells allowed us to confidently identify Cx40^+^ single artery ECs and conclude that at least a sub-population of artery ECs, upon MI, express Apj↓an indicative of vascular de-differentiation.

The de-differentiation indicators were absent in adults. This finding positively correlated with the ability of artery ECs to re-enter cell cycle. Neonates had much higher number of cycling artery ECs than P7 or adults. This finding was verified via two independent methods—single cell gene expression analyses of cyc-aECs and presence of *Cx40CreER*^+^ EdU^+^ cells in an *in vivo* cardiac MI model. Thus, lack of de-differentiation limits the number of cycling aECs in non-regenerative hearts and consequently affects their ability to form collateral arteries by Artery Reassembly.

### Reactivation of Developmental Pathways upon Injury?

Several molecular pathways relevant to vascular development regulate EC response to injury^65^. For instance, Notch^66^, Apelin^45^, Cxcl12^67–70^ signaling, in mice, regulate different aspects of vascular development and remodeling during cardiac morphogenesis; ranging from EC fate determination to regulation of different EC responses such as migration, proliferation and tube formation. Interestingly, some of these pathways are also associated with EC regeneration upon injury^1,24,48^, making them potential therapeutic candidates for treating vascular conditions in human patients^71^. Further studies are required, and will elucidate if (and how) the cellular and molecular mechanisms associated with these signaling pathways, during development and injury responses, are similar.

Vegf signaling drives coronary development in zebrafish and mouse hearts. In mice, Vegf signaling drives migration of coronary endothelial^43,72^ and endocardial endothelial cells^58^ in developing hearts, and proliferation of coronary ECs in heart explant cultures^73^. In zebrafish, *vegfa* mutants show reduced EC proliferation possibly via PI3K signaling^74^. Upon cardiac injury, endocardial Vegfa signaling^1^ and Vegfc-mediated (Cxcl8a/Cxcr1-driven) coronary EC proliferation^75^ execute revascularization and cardiac regeneration of ischemic hearts. In this study, we highlight the role of VegfR2-mediated signaling in artery EC response triggered by neonatal cardiac injury. We show that VegfR2 in neonates is expressed in coronary artery ECs, deletion of which, significantly reduced the number of collateral arteries formed via Artery Reassembly.

Our data show that artery tips do not proliferate. Proliferation is rather demonstrated by single aECs which originate from these artery tips. This suggests that migration of pre-existing artery ECs could precede artery EC proliferation in the watershed. Appearance of *VegfR2* depleted single artery ECs in ischemic watershed regions suggested that artery EC migration was not affected. VegfR2 depleted single artery ECs showed reduced proliferation potential, which could account for lack of lineage-traced collateral arteries in the knock out hearts. Proliferation of artery ECs is followed by their coalescence into collateral arteries. Thus, we cannot exclude the possible role of Vegf in coalescence of artery cells, or the vascular steps that follow, like lumenization or maturation of collateral arteries.

The precise source of Vegf ligand in injured mouse hearts could be the endocardium^76^, myocardium (Vegfa)^58,77^, epicardium (Vegfc^43^, or Vegfa^78^), or cardiac endothelial cells^75^, thus suggesting an autocrine or paracrine signaling as observed in injured zebrafish hearts^75^. In our neonatal mouse single cell dataset, we found both *Vegfa* and *Vegfc* being expressed by mature artery ECs (aEC1-4), but at lower levels by cyc-aECs (or by trans-ECs, SMCs, endoMT ECs, endo ECs), indicating a possible cell autonomous mechanism. However, an alternate source of Vegf ligand, such as, cardiomyocytes, fibroblasts or immune cells, which are excluded from our dataset, cannot be neglected.

### Role of Vegf Pathway in Human Heart Disease and its Treatment

Single nucleotide polymorphisms (SNPs) in both introns and exons of *KDR* (murine *VegfR2*) are strongly associated with coronary heart disease (CHD)^79^. These SNPs can either modulate the affinity to ligand, or alter E2F binding to *KDR* promoter, subsequently, affecting *KDR* activity and downstream signaling^79^. Elevated levels of circulating VEGF ligands are positively and significantly associated with CHD^80^, coronary artery disease (CAD)^81,82^ and coronary artery lesions^83,84^. Several SNPs affect VEGF levels^80,82^ and may account for the variation observed in revascularization abilities in individuals across different genetic backgrounds. At least three such SNPs in *VEGFA* are strongly associated with increased risk of CHD/MI^85^ and one with coronary collateral vessel perfusion^86^. How the association of VEGF/KDR signaling axis with CHD may serve for better prognosis or indicate the need for vascularization, remains unclear.

Preclinical studies in animals with sparse^87^ to no^54^ pre-existing collateral arteries have consistently elucidated role of Vegf in ischemia-induced collateral formation. On the contrary, numerous past and ongoing clinical trials with *VEGF* alone, or combined with pro-angiogenic/ regenerative factors, have been concluded with little to no success (data available at clinicaltrials.gov)^87,88^. The delivery methods appear to be safe, but the angiogenic effects, particularly the efficacy of *VEGF* gene therapies in stimulating reperfusion via collateral arteries is minimal^87^. Mechanistic details of Vegf’s mode of action, in orchestrating vascular cell responses that lead to collateral formation, could aid designing clinical trials with higher efficacies.

We show that mouse coronary ECs in MI hearts, de-differentiate and then expand in VegfR2-dependent manner. The mere presence of a significant number of single artery ECs in a limited watershed area could be sufficient to create new arteries^31^. Thus, it will be interesting to see if this threshold density of artery ECs is accomplished by Vegf-mediated proliferation. Similarly, identifying the molecular drivers of EC de-differentiation is also crucial, and when combined with Vegf, could promote collaterals in individuals with CHD and CAD.

## Authors Contribution

SD and KR conceived the idea. SD, KR, YJW provided resources. GA performed experiments with neonatal MI model. Sneha K and Suraj K performed bioinformatics analyses. SD, GA and Sneha K wrote the manuscript. HW performed adult MI surgeries. PERC and BB performed some neonatal MI surgeries. SD and KMG performed FACS.

## Acknowledgements

We thank the following facilities for their technical support: Stanford Genome Sequencing Center, Stanford shared FACS facility, Animal Care and Resource Center at NCBS, Mouse Genome Engineering Facility at NCBS, Central Imaging and Flow Cytometry Facility at NCBS, Dr. Dhandapany Perundurai’s lab from Institute For Stem Cell Science and Regenerative Medicine (InStem), for their technical help with echocardiography.

## Data availability

Raw sequence files and Cell Ranger processed data are available on GEO (accession ID: GSE210307)

Scripts used for the analysis are available at https://github.com/Snehasrivatsa/scRNA

## Methods

### Tissue digestion and single cell sorting

Neonatal and adult mouse hearts were dissected and atria was removed from each heart. Tissue digestion was performed only with cardiac ventricles. Adult mouse hearts were additionally perfused before harvesting hearts. ∼15-20 neonatal (for sham and MI each) and ∼10-12 adult (for sham and MI each) *Cx40CreER*; *Rosa26^TdTomato^* mice were used for tissue digestion. Post-explant, ventricles of the hearts were minced using razor blade and fine forceps and transferred to a tissue digestion buffer. The tissue digestion buffer was composed of 500U/ml Collagenase IV (Worthington, Catalog# LS004186), 1.2U/ml Dispase (Worthington, Catalog# LS02100), 32U/ml DNase I (Worthington, Catalog# LS002007) in commercial grade sterile 1X DPBS with Ca^2+^ and Mg^2+^. Each heart was transferred to ∼300µl of digestion buffer. Hearts bathing in digestion buffer were incubated in 37° Celsius shaker for 40-45 minutes, with periodic resuspension using micropipettes to break down tissue lumps. Digested tissue was then equilibrated with 4 volumes of 5% FBS prepared in 1XDPBS, gently mixed with a micropipette and filtered using 40µm of sterile cell strainer. The filtered tissue digest was spun down for 5 minutes at 400g at 4° Celsius, and the cell pellet was gently dissolved in 5ml of 3% FBS in 1XDPBS. The re-suspended cell pellet was spun down for 5 minutes at 400g at 4° Celsius. Supernatant was discarded and the cell pellet was suspended in ∼500-600µl of 3% FBS in 1XDPBS. These suspended single cells were stained with Ter119-Alexafluor-647 (Biolegend, Catalog# 116218, 1:100 dilution) for 30 minutes at 4° Celsius. Post-staining, cells were washed with and re-suspended in fresh 3% FBS/ 1XDPBS, followed by single cell sorting with fluorescent activated cell sorter. DAPI was added prior to starting the cell sort. Live single endothelial cells (DAPI^−^ Ter119^−^ TdTomato^+^) were collected, in that order, using BD FACS Aria II cell sorter. Single cells were counted, and submitted to Stanford Genome Sequencing Service Center for 10X single cell V3 library preparation. Sequencing was performed using HiSeq 4000.

### Single Cell RNA sequencing data analyses

#### Processing of FASTA files

Illumina reads obtained were processed on Cell Ranger (10X Genomics). To demultiplex the BCL files and convert them to FASTQ files, the *mkfastq* function was used. The reads were aligned to mm10 (mouse genome) and TdTomato sequences with the count function.

#### Preprocessing on R

The output files of Cell Ranger were imported into RStudio v4.05 using Seurat v4.03. This was performed with *Read10x* feature by providing path to a folder with zip files of expression matrix, features and barcodes files. The cells were analyzed for nFeature RNA count and percent of mitochondrial RNA using the standard pre-processing mentioned on the Seurat^28^. The tutorial is available at https://satijalab.org/seurat/articles/get_started.html.

The percentage of mito genes expressed in the cells of our dataset were less than 1, hence no cells were removed. Based on Q/C violin plot, all cells with >250 nFeature RNA count were analyzed. These cutoffs were determined from quality control analyses of all datasets, which include, subsets of cardiac cells filtered with expression of *Cadherin5*, *Pecam1* or TdTomato from uninjured and injured neonates and adults (Figure **S1A**, **S2A**, **S9A, E**). We obtained and analyzed the following number of cells from each subset. *Cadherin5* >1: 5432 sham and 2200 MI neonatal cells, *Cadherin5*>1: 6256 sham and 1690 MI adult cells, *Pecam1*>1: 5520 sham and 2152 MI neonatal cells, *Pecam1*>1: 6228 sham and 1774 MI adult cells, TdTomato>0: 5557 sham and 497 MI neonatal cells, TdTomato>0: 3090 sham and 892 MI adult cells.

DoubletFinder, a package on RStudio was used to check if there were any doublets incorporated in the dataset. The analysis showed that, 7.5% of all cells were doublets in both sham and MI datasets. These doublets were mostly present in endoEC, endoMT and SMC cell clusters which were excluded from the study. Trans-EC, a unique cluster expressing marker genes for multiple cell types was considered specifically for the analysis (as the chance of it being a doublet is higher). No doublets were found in MI trans-EC cluster. In the sham trans-EC cluster, there were 3 doublets.

#### Normalization, scaling and clustering

Data was log normalized with scale factor set to 10000, followed by finding variable features with n set to 2000 and finally, scaled with *scaledata*. We focused on artery endothelial cells, by sub-setting the data using different markers. *Cadherin5* or *Pecam1* (pan endothelial markers) expression greater than 1 were analyzed. A filter for greater than 1 was used for *Cadherin5* or *Pecam1*to exclude non-ECs expressing these genes at a minimal level. As the cells from both filters echoed the same clustering pattern, we used *Cadherin5* cells for further analyses. The pattern acquired by clustering sham and MI data separately or together (merged) echoed the same patterns. Hence, the sham and MI data were integrated using the *IntegrateData* feature from the Seurat package. Normalization, finding variable features and scaling data was repeated on the integrated data, using the same settings as above.

We performed Principal Component Analysis (PCA) using the top 30 PCs, from integrated data. We did not observe any variations in clustering patterns when 10-100 PCs were used. Clusters for the integrated data were obtained with Louvain algorithm with resolution set to 0.5 (https://satijalab.org/seurat/articles/pbmc3k_tutorial.html). The cells were projected and visualized in two-dimension using Uniform Manifold Approximation and Projection for Dimension Reduction (UMAP); parameters were n.neighbors set to 10, dims set to 1:20, spread set to 2, and min.dist set to 0.3. Attributes *group.by* and *split.by* were used to observe sham and MI cells. This was done for optimal spread of cells and best visualization.

TdTomato sequence was aligned with and checked in our datasets. Thus, cells with TdTomato expression greater than 0, were analyzed. Since the MI cell number was 10 fold lesser than that of the sham, data was analyzed by sub-setting the merged data based on its orig.ident and re-clustering the respective cell populations. The steps used with *Cadherin* >1 cells were used for all clustering steps with TdTomato >0.

For expression of genes, data was sub-setted for gene >0, the cell numbers per cluster were obtained using table.

#### Cluster identification

The communities detected were identified with Violin plots (*StackedVlnPlot* from CellChat v1.1.0 package (https://github.com/sqjin/CellChat), *DotPlots* (a feature of Seurat package to visualize gene expression), and feature plots (Seurat argument). Clusters were identified with established markers for a variety of cardiac cell specific genes. Cluster identifications were confirmed using *AddModuleScore* feature from Seurat package. A list of cell-type specific, established markers were used for this analysis on *Cadherin5*>1, *Pecam1*>1 and TdTomato>0 datasets in both neonate and adult datasets. Average expression of the listed genes for each cell type was visualized using a feature plot. Neonatal trans-EC cluster did not show unique expression of any known cardiac cell types.

Heatmaps were used to check the variation in expression of de-differentiation genes across sham and MI datasets. *DoHeatmap* feature from Seurat was used to plot the genes of interest. Violin plots generated from Seurat package were analyzed by using the *split* attribute.

#### Cell cycle analysis

Updated G2M and S phase specific gene lists (from 2019) on Seurat were used. As the genes were in human gene format, *useMart* function from biomaRt v3.12 (RStudio package) was employed to obtain hsapiens_gene_ensembl and mmusculus_gene_ensembl from ensembl database. Homology mapping and gene symbol conversion of genes from 2 gene-sets was performed using *getLDS* function. Genes were converted to mouse gene format using *convertHumanGeneList*. Cell cycle analysis was performed using standard cell cycle regression (CellCycleScoring) on Seurat.

#### Trajectory analysis

Monocle3^89^ was used for trajectory analysis to understand possible pseudo time relationship between the cell types. Seurat object split into sham and MI from the integrated data, subsetted for ECs alone, was used for the analysis. *as.cell_data_set* feature from SeuratWrappers was used to convert Seurat object to Monocle3 object. Further, the Monocle3 object was subjected to pseudo time analysis using *learn_graph* and *order_cells* functions (https://cole-trapnell-lab.github.io/monocle3/docs/starting/). Results were visualized on UMAPs with the aid of *plot_cells* function.

#### RNA Velocity

Spliced and unspliced counts were generated from raw Fastq files using kallisto bustools wrapper *kb* (kb-python v0:27:0, https://www.kallistobus.tools/)^90^. Mouse genomic (DNA) FASTA and GTF annotations were downloaded from Ensembl and a transcriptome index was generated by kallisto using *kb ref* command. *Kb count* command was used to generate the count matrix in the form of h5ad file. The h5ad file was converted into a seurat object using SeuratDisk (v0.0.0.9019). The preprocessing, normalization, scaling and clustering were done as described earlier. The RNA velocity maps were generated using Velocyto package (velocyto.R v6)^91^.

#### Analysis of differentiation state

R package CytoTRACE v0.3.3^53^ (https://cytotrace.stanford.edu/) was used for cell differentiation analysis. Cell counts and UMAP embeddings were obtained as data frames from sham and MI clusters separately, for the analysis. A CytoTRACE score was assigned to each cell with the CytoTRACE function, which was visualized on the UMAP embeddings with *plotCytoTRACE* feature. We compared the mean CytoTRACE scores between all EC clusters. To do so, CytoTRACE score for each cell and its cluster number was obtained. The mean CytoTRACE score was then calculated for each cluster. These mean values along with standard deviations were plotted as boxplots using *ggboxplot* feature from ggpubr v0.4.0 package.

#### Gene Ontology (GO) analysis

List from *FindAllMarkers* was used to recognize processes that appear on publicly available online tool GOrilla (Gene Ontology enRIchment anaLysis and visuaLizAtion tool)^92^ (http://cbl-gorilla.cs.technion.ac.il/). Gene Ontology analyses was performed on MI gene list, with sham genes serving as background. Default settings with a filter of p-value less than 10^−3^ were used. Processes with significant p-values (<0.0005), and relevant to our study, were plotted on a bar graph.

#### g:Profiler

Pathway analysis was performed using an online tool g:Profiler^93^. The gene list was obtained for neonatal cluster aEC1 (from merged *Cadherin5* >1 dataset), using *FindMarkers* feature, with test.use set to “MAST”. The organism was set to *Mus musculus* and the pathways were obtained from databases like KEGG, reactome, WikiPathways etc.

#### Statistical analysis on sequenced data

Wilcoxon test was performed using *RunPresto* feature from SeuratWrappers package. The statistical test was performed on cyc-aEC and aEC1 cells from neonatal *Cadherin5*^+^ MI group. Additionally, the test was performed on *Cadherin5*^+^ cyc-aECs from neonatal and adult MI datasets. For this, the data was merged using the *merge* feature from Seurat, and retained as such, without any further modifications. Test was done to check statistical significance for expression of *Cx40*, *Cxcr4*, *Aplnr*, *Car4*, *Apln*, *Foxm1, Vegfa*, *Vegfc*, *Kdr* across different treatments (sham and MI in Figure **4**) or clusters (cyc-aEC and aEC1 in neonates, in Figure **S7A**).

#### Experimental model

Following mouse lines were used in this study. *Rosa26^TdTomato^* Cre reporter line (The Jackson Laboratory, B6.Cg-Gt(ROSA)26Sortm9(CAG-TdTomato)Hze/J, Stock# 007909), *Cx40CreER*^94^, *VegfR2 flox* (The Jackson Laboratory, *Kdr^tm2Sato^*/J, Stock#018977), *Cx40^eGFP/+^*^96^.

All mice were housed and bred either in Stanford (in accordance with institutional animal care and use committee (IACUC)) or NCBS animal facility (in accordance with Institutional Animal Ethics Committee (IAEC)). All Tamoxifen injections (6mg dissolved in corn oil) were administered intraperitoneally to nursing mothers for Cre activation in new born pups. Efficiency of Tamoxifen and Cre activation has been shown earlier to be ∼84% using Cx40CreER mice and ∼86% using ApjCreER mice^24^. Data shown in this study is compiled from age-matched males and females from several litters.

#### Experimental study design

Sample size for the current experiment was determined from our previous study conducted with the same animal model with similar treatments^24^. In any given litter, animals were categorized as control or knockout based on their genotype, and irrespective of the sex. Severity of MI was determined by first checking the location of LCA ligation under fluorescence stereo-microscope (to check knot around TdTomato^+^ arteries). Hearts with severe MI (where either the main LCA or the two primary branches of LCA were captured), were included in this study. Hearts with mild MI (where secondary, or tertiary branches were ligated), were excluded from the study. Biological replicates from multiple litters were included in the study, and are mentioned below for each in vivo experiment.

#### Genotyping information

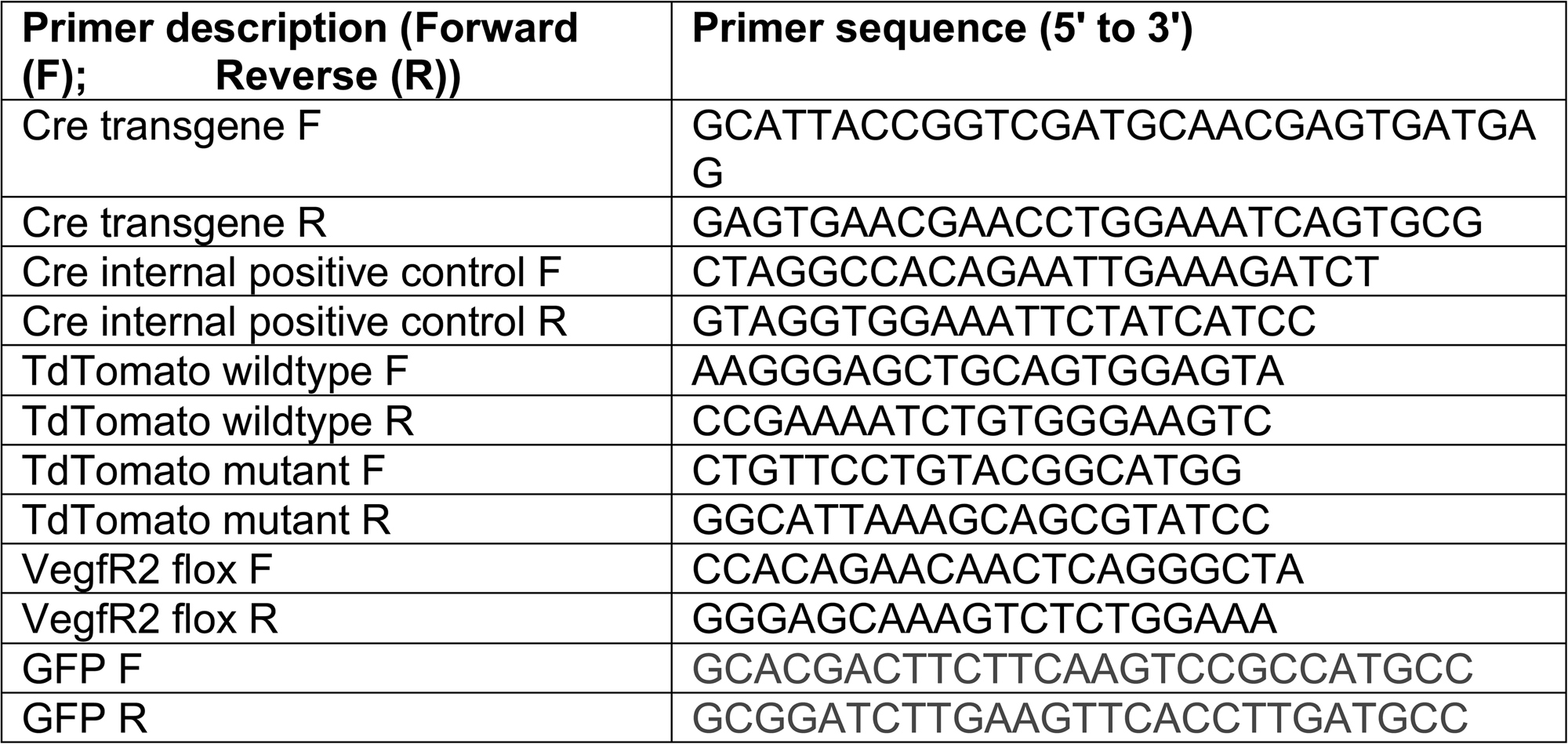

#### Adult LCA ligations

Adult mice were anesthetized in a 4% isoflurane chamber and then intubated with a 20G angiocatheter. General anesthesia was maintained with 1.5-4% isoflurane during mechanical ventilation at a respiratory rate of 180 breaths per minute. The mice were placed in right lateral decubitus position and the left chest was shaved and sterilely prepped. 1 mg/kg Bupivicaine and 0.5 mg/kg Buprenorphine were injected subcutaneously for perioperative analgesia. A 1 cm left thoracotomy incision was made, and the pericardium was opened. The left coronary artery was identified and permanently ligated using a 6-0 polypropylene suture. The chest was then closed in layers using 5-0 polypropylene sutures. Isoflurane was then weaned off and the mice were extubated after regaining physiological respirations. 5mg/kg Carprofen was injected subcutaneously daily for the first three days after surgery for postoperative analgesia. For adult MI experiments, 2-3 months old mice were used (both males and females). To activate Cre, a single dose of 6mg Tamoxifen intraperitoneal injections were administered to each mouse, 2 days prior to MI. Echocardiograms were obtained from P32/P33 adult mice (sham or MI surgeries at P2) ∼30days post-MI using VisualSonics Vevo 3100 imaging system (FUJIFILM VisualSonics Inc.) with an MX400 20–46 MHz transducer. Data was analyzed using Vevo LAB version 3.1.0 (Build 13029)

#### Neonatal LCA ligations

Technical details of the procedure are referred and modified from Mahmoud et al^95^. To induce hypothermic arrest, neonatal mice (P2 or P7) were incubated on ice. Neonates were wrapped with a 3-layer, pre-cooled, gauze piece for ∼3 minutes. Anesthesia was confirmed using toe pinch. Surgery was performed in sterile conditions from start to end, using sterile surgical tools at all times. Animals were placed in a supine position, on a sterile surgical drape, wrapped around cold ice pack. Animals were then prepped with betadine and 70% ethanol. Left anterolateral thoracotomy was performed by making an incision on fourth intercostal space to access heart, under a dissecting microscope. The left coronary artery (LCA) was identified and ligated by passing a needle attached to 8-0 non-absorbable prolene suture (Ethicon Ethilon, catalog# NW3322). Post-ligation, the 8-0 prolene suture was used to close the ribs, intercostal muscles and skin separately. The neonate was then transferred to warm plate (37° Celsius) to gain consciousness and recover completely. When conscious, neonates were returned to their mother’s cage and closely observed over the next few days.

#### Whole mount immunostaining of neonatal hearts

The procedure has been described earlier^24^. Neonatal whole hearts were dissected and immediately fixed in 4% paraformaldehyde at 4° Celsius for 1hr on a rocker. Hearts were then subjected to three, 15 minutes washes, with 1X PBS at 4° Celsius. Primary antibody staining was done by incubating hearts in respective dilutions of primary antibodies, prepared in at least 5 volumes of 1X PBS containing 0.5% Triton-X 100 (0.5% PBT). Incubations were done overnight at 4° Celsius, on a shaker. Hearts were washed in 20 volumes of 0.5% PBT for 12 hours, with a change in wash buffer (0.5% PBT), every 2 hours at 4° Celsius. The secondary antibody staining was performed by incubating hearts in 1:250 dilution prepared in 0.5% PBT. Incubation was done on a shaker. Next, hearts were washed in 20 volumes of 0.5% PBT, for 4 days with a change in wash buffer, every two hours at 4° Celsius. All washes were performed with continuous but gentle shaking. Hearts were cleared in at least 2 volumes of Anti-fade mounting media (https://www.jacksonimmuno.com/technical/products/protocols/anti-fade) for 2 hours, in dark, at room temperature. Whole hearts were then stored at −20° Celsius till imaged. Post staining washes were done in 15 and 50 ml tubes while incubation in antibodies or mounting media were performed in 1.5ml microtubes.

Neonates were injected with 30µl of 10mM 5-ethynyl-2’-deoxyuridine (EdU) (prepared in 1X commercial grade PBS, catalog #10010023) for 1hour, prior to harvesting their hearts. EdU label detection was combined with post-secondary immunostaining steps as described above, and, detected using Click-iT staining kit (Invitrogen, Catalog# C10340).

#### Confocal imaging of neonatal hearts

Whole neonatal (immunostained) hearts were mounted on a double concave microscope slide (Sail brand, catalog# 7104) using a microscope coverslip (0.13-0.17mm, Thermo Scientific, Catalog# 48367-059) so that the ventral watershed regions spanning left and right coronary arteries can be imaged. Imaging was performed using Olympus FV3000 upright confocal microscope. Whole hearts were imaged using 4X, 10X, 20X or 60X (oil) objectives. Multiple z stacks were acquired using FV31S-SW software. Multiple images of the whole heart were processed on Fiji-ImageJ software, and stitched with Adobe Photoshop.

#### Antibodies

Primary antibodies: Anti-CX40 (1:500; Alpha Diagnostics Int. Inc.; Catalog# CX40-A, RRID: AB_1616128), Anti-VEGFR2 (1:125, R&D Systems, Catalog# AF644, RRID: AB_355500). Secondary antibodies: Alexa fluor-conjugated antibodies (488, 555, 633) from Invitrogen were used at 1:250 dilutions.

### Quantification from images

#### Collateral number

Number of *Cx40CreER* lineage-traced (TdTomato positive) collateral arteries were quantified from images acquired using 10X objective from whole hearts. Number of stacks (12-14) and step size (2µm) of the images used, were consistent across genotypes. Data was compiled from 6-9 litters from each genotype. 8 control, 5 heterozygous and 14 knockout hearts were used for quantification. An unpaired t-test was performed to check statistical significance.

#### Cx40 intensity in proliferating artery ECs

Proliferating TdTomato^+^ Cx40^+^ cardiac artery ECs (4 days post-MI) were identified with EdU^+^ nuclear labelling. Cytoplasmic intensity of Cx40 in these (TdTomato^+^ Cx40^+^ EdU^+^) ECs were quantified using images acquired with 60X (oil) objective. Mean fluorescent intensity (MFI) of cytoplasmic Cx40 signal was measured using a selection tool available with Fiji-ImageJ software. Quantification was performed with 68 single TdTomato^+^ Cx40^+^ EdU^+^ artery ECs, found in the watershed of 4 hearts, collected 4 days post-MI. Comparison between proliferative (EdU^+^) and non-proliferative (EdU^−^) artery ECs were made within the same field of view. An unpaired t-test was done to check statistical significance. Similar analysis was done on 67 artery ECs within artery tips or integrated into mature arteries, as identified by Cx40 immunostaining. Data was compiled from 5 MI hearts from different 4 litters.

#### Single artery EC proliferation in control and *VegfR2* knockout hearts

EdU^+^ TdTomato^+^ single artery ECs in the watershed regions, were quantified using images acquired with 20X objective. On an average 20-30 TdTomato^+^ single artery ECs were observed per watershed (each field of view), which was consistent across genotypes. Quantification was performed on 6 control (36 field of views) and 5 *VegfR2* knockout (24 field of views) hearts. A total of 784 control TdTomato^+^ artery ECs and 512 *VegfR2*knockout TdTomato^+^ artery ECs were counted for this quantification. Data was compiled from 3-5 litters from each genotype. 6 control and 5 knockout hearts were used for quantification. An unpaired t-test was performed to check statistical significance.

#### Single artery EC proliferation in P11 MI hearts (MI at P7)

EdU^+^ TdTomato^+^ proliferating artery ECs were counted in artery tips and saECs from 3 *Cx40CreER*; *Rosa26^TdTomato^* hearts. ∼80-120µm of distal ends of arteries were identified as artery tips. 25 fields of view from P11 watershed regions, from images taken with 60X (oil) objective, were used to count a total of 113 TdTomato^+^ residual single cells.

#### TdTomato fluorescence in control and *VegfR2* knockout watershed

Mean Fluorescence Intensity was measured using Fiji for *Cx40CreER* lineage-traced TdTomato^+^ single artery ECs found in watershed area 4 days post-MI. Images of watershed regions (each field of view) lacking any large or collateral arteries were acquired with 20X objective. Watershed images only with single artery cells were considered for this quantification. 3 hearts from each genotype were used for this quantification. An unpaired t-test was performed to check statistical significance.

#### LCA width

LCA width was measured using a line selection tool on Fiji. The analysis was done for 3 control and 2 *VegfR2* KO hearts. Width was measured for the main branch of LCA in all hearts and genotypes. An unpaired t-test was performed to check statistical significance.

## Supplemental Figures and legends

**Supplemental Figure 1:**
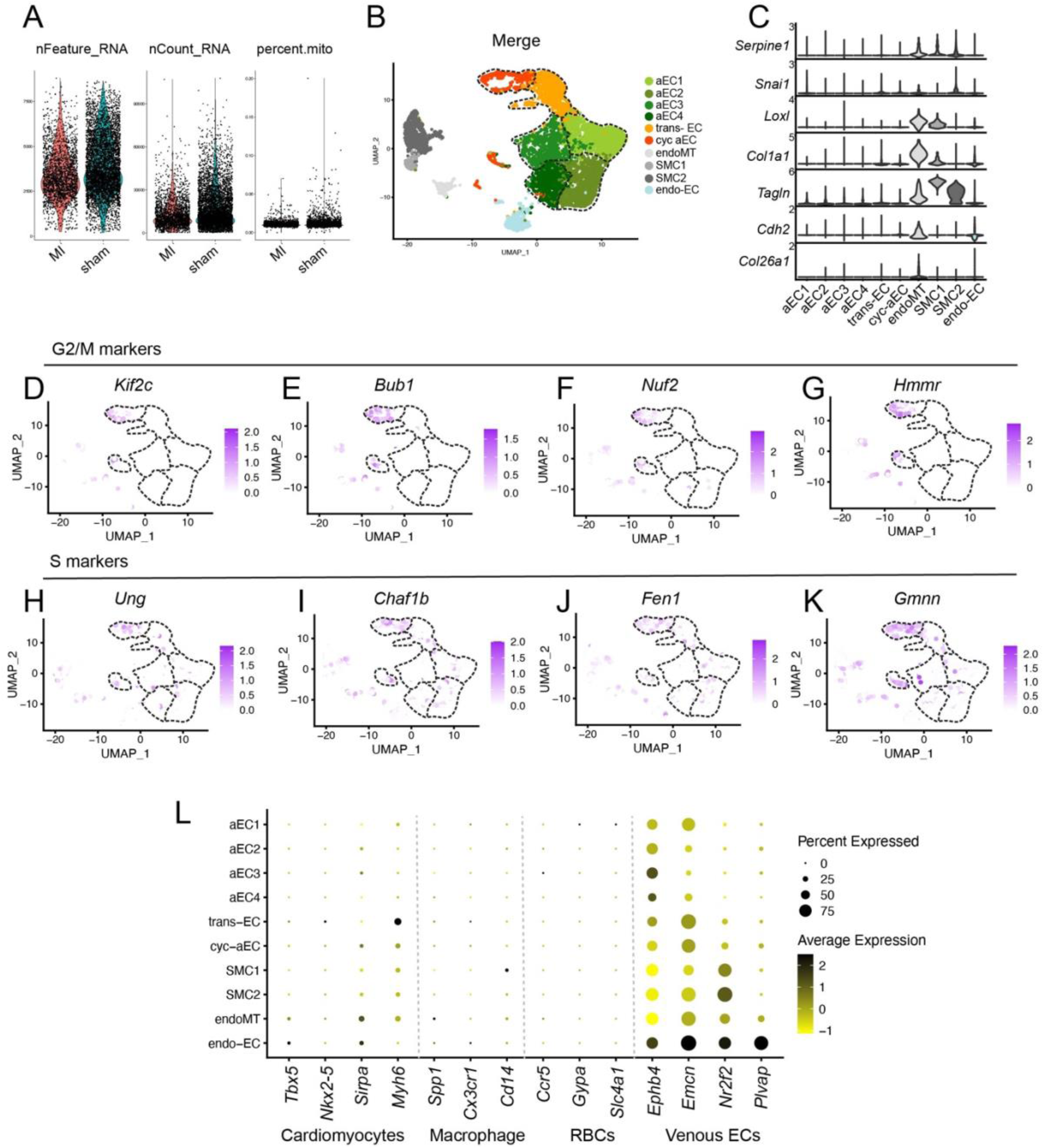
Identification of neonatal *Cadherin5*^+^ cell clusters. (**A**) Violin plots showing quality control (analyses of nFeature counts, nCount_RNA and mitochondrial genes) performed on neonatal *Cadherin5*^+^ dataset. (**B**) Visualization of different identified cell clusters, in neonatal *Cadherin5*^+^ dataset, using UMAP. (**C**) Violin Plot showing expression of endo-MT genes in neonatal *Cadherin5*^+^ cells. (**D**-**G**) Feature plots showing the expression of genes specific to G2/M phase. (**H**-**K**) Feature plots showing the expression of S phase markers in neonatal *Cadherin5*^+^ cells. (**L**) Dot plot showing parentage of cells and average expression of genes, specific to different cell-types.

**Supplemental Figure 2:**
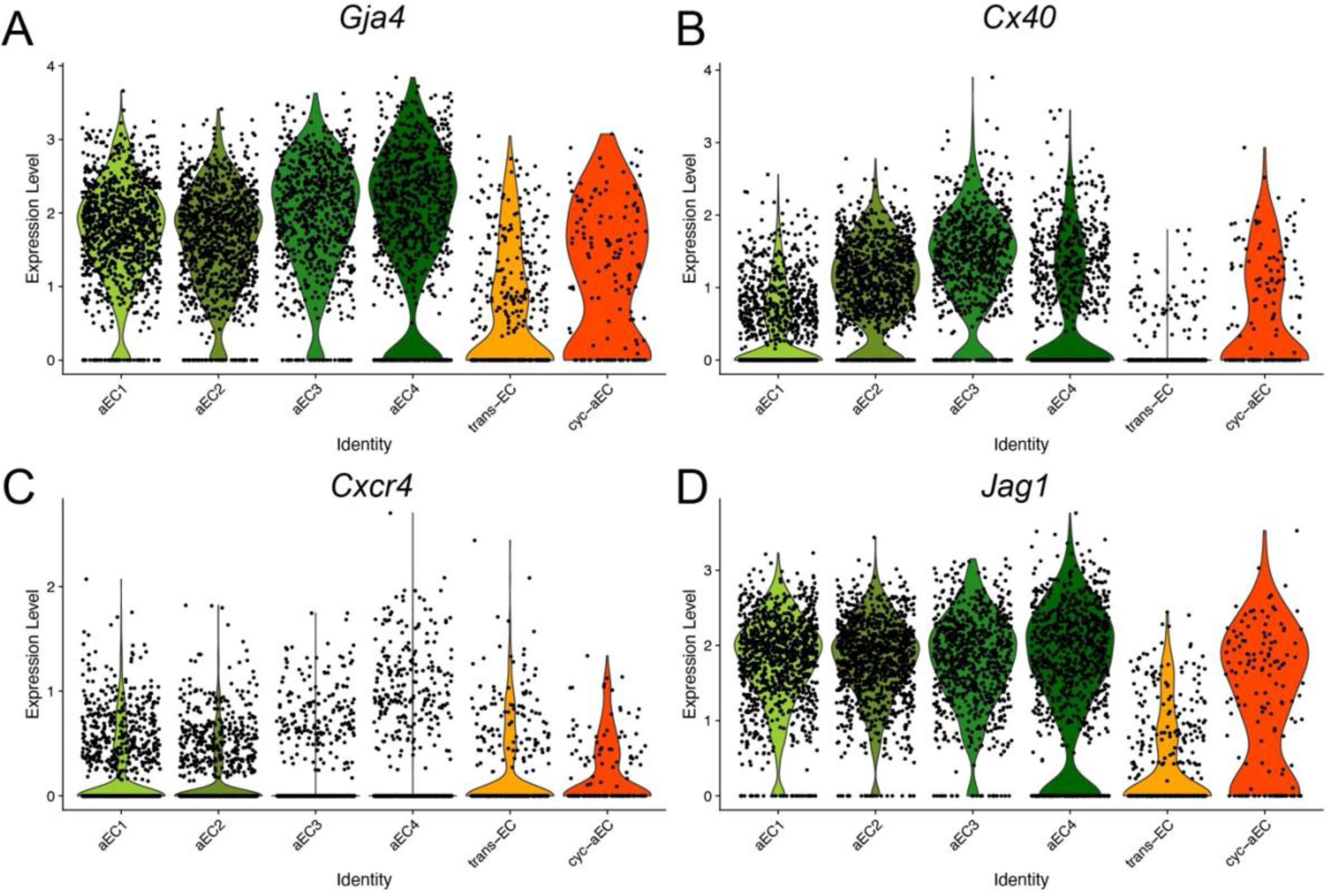
Artery specific gene expression in neonatal *Cadherin5*^+^ cycling aEC cluster. (**A**-**D**) Violin plots showing the expression levels of artery endothelial cell specific markers in *Cadherin5*^+^ neonatal sham cells.

**Supplemental Figure 3:**
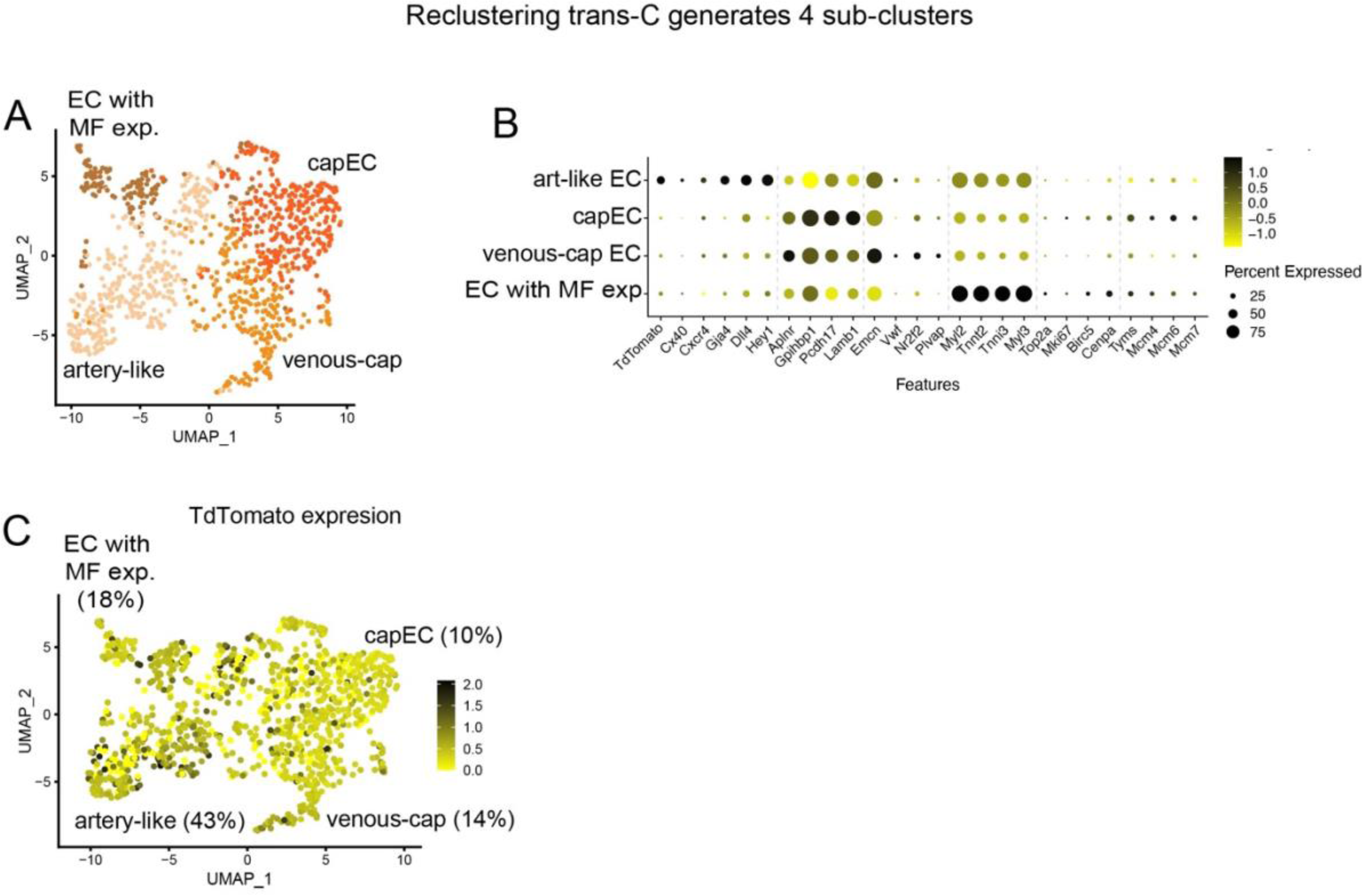
Gene expression analyses of neonatal *Cadherin5*^+^ trans-EC cluster. (**A**) Visualization of cells upon re-clustering trans-EC cluster from neonatal dataset, using UMAP. (**B**) Dot plot showing average gene expression and percentage of cells expressing various genes expressed in ECs. (**C**) Feature plot showing distribution of TdTomato expressing cells in **A**. MF, myofibrillar; cap, capillary; exp, expression.

**Supplemental Figure 4:**
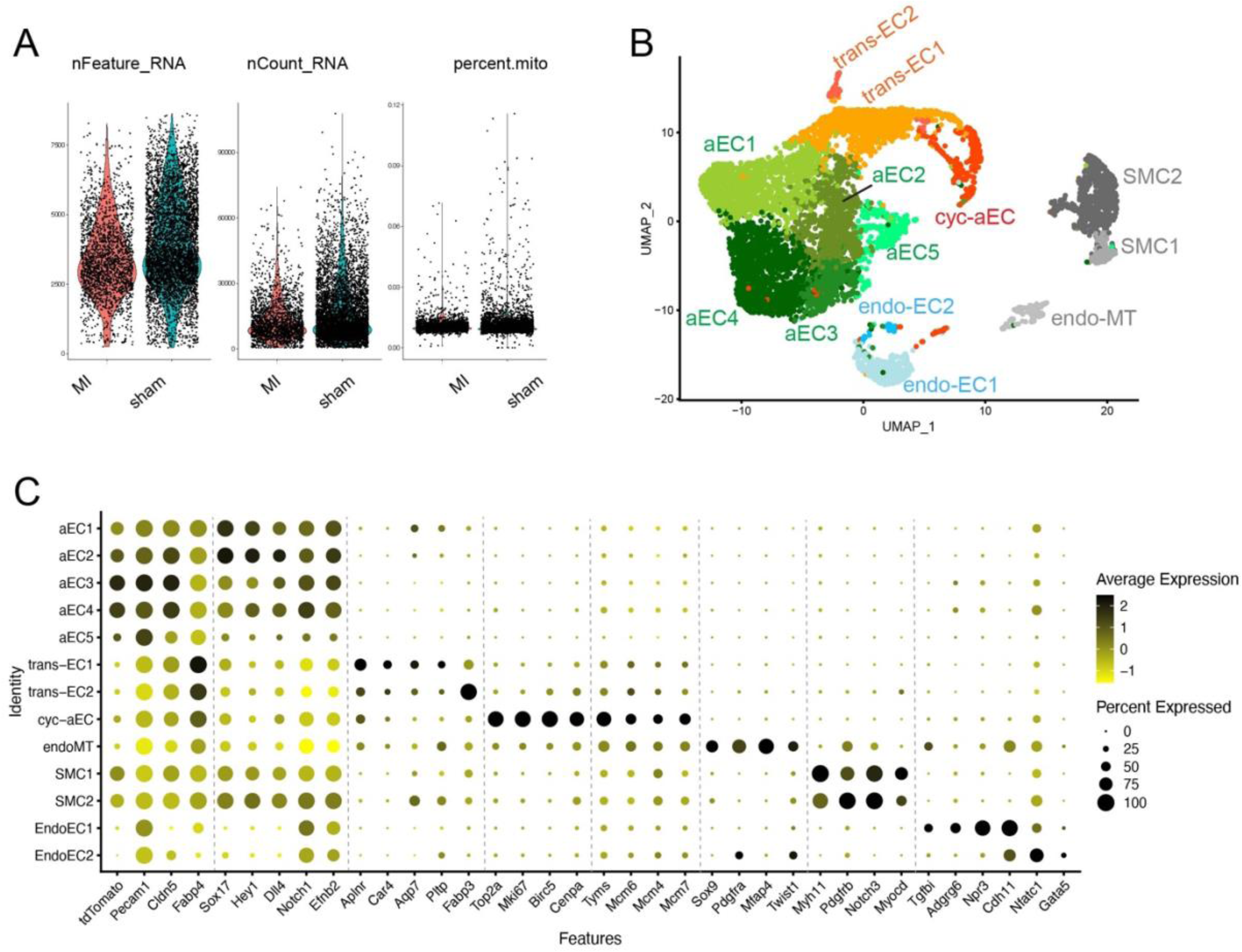
Identification of neonatal *Pecam1*^+^ cell clusters. (**A**) Violin plots showing quality control (analyses of nFeature counts, nCount_RNA and mitochondrial genes) performed on neonatal *Pecam1*^+^ dataset. (**B**) Visualization of clusters in neonatal *Pecam1*^+^ cells, using UMAP. (**C**) DotPlot showing percentage of cells and average gene expression of cell-type specific genes.

**Supplemental Figure 5:**
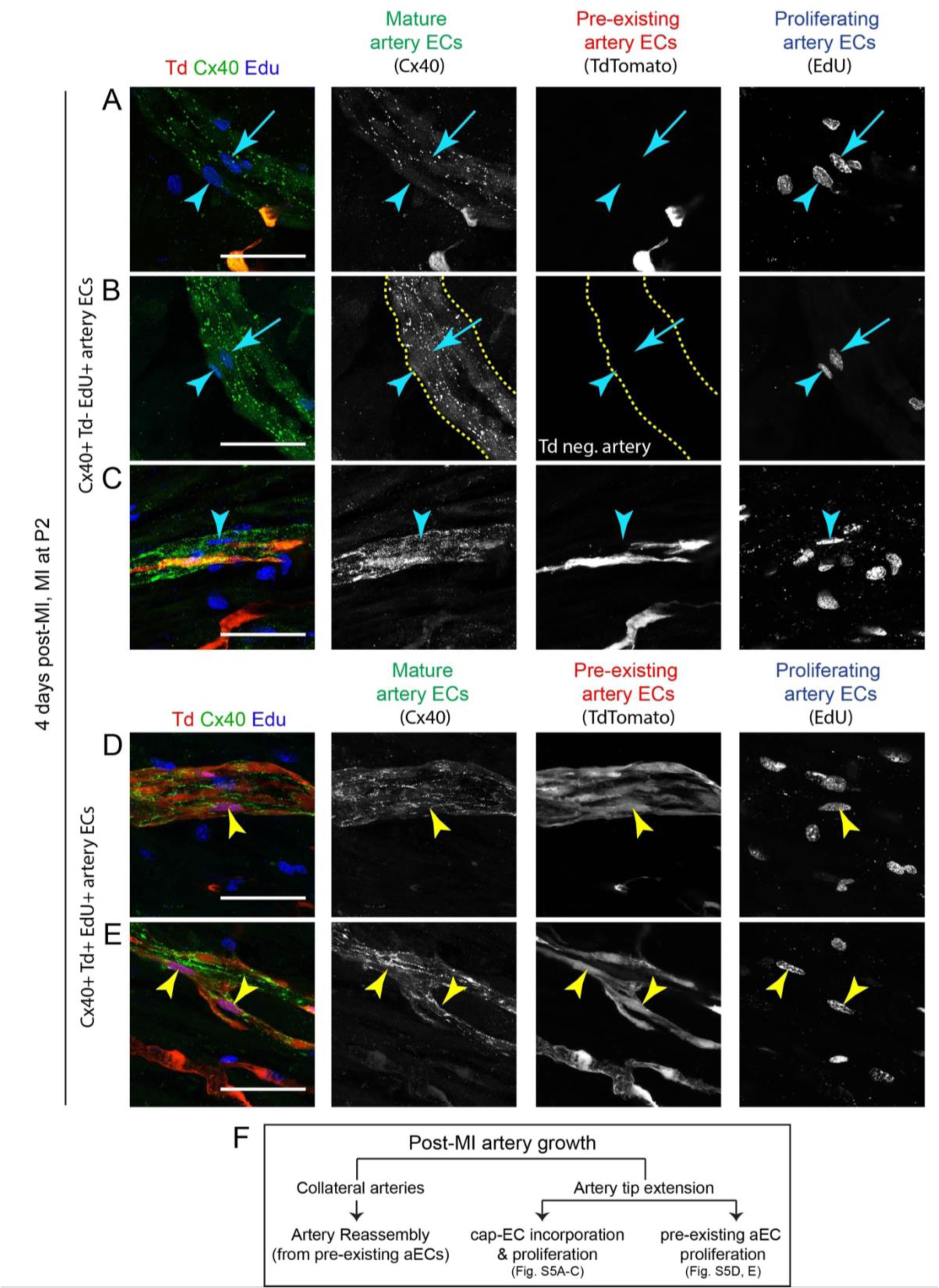
Analyses of post-MI artery EC proliferation in neonatal hearts. Representative confocal images of artery segments from neonatal P6 hearts, 4 days post MI, immunostained for Cx40, lineage traced for *Cx40creER* (with TdTomato) and labelled for EdU. Images are taken from ischemic regions (watershed area below the ligation). Arrows and arrowheads show (**A**-**C**) EdU^+^ Cx40^+^ Tdtomato^−^ artery ECs and (**D**, **E**) EdU^+^ Cx40^+^ Tdtomato^+^ artery ECs. (**F**) Modes of post-MI artery growth in neonatal hearts. neg., negative; aEC, artery endothelial cells; cap-EC, capillary endothelial cells. Td, TdTomato. Scale bar: 50μm

**Supplemental Figure 6:**
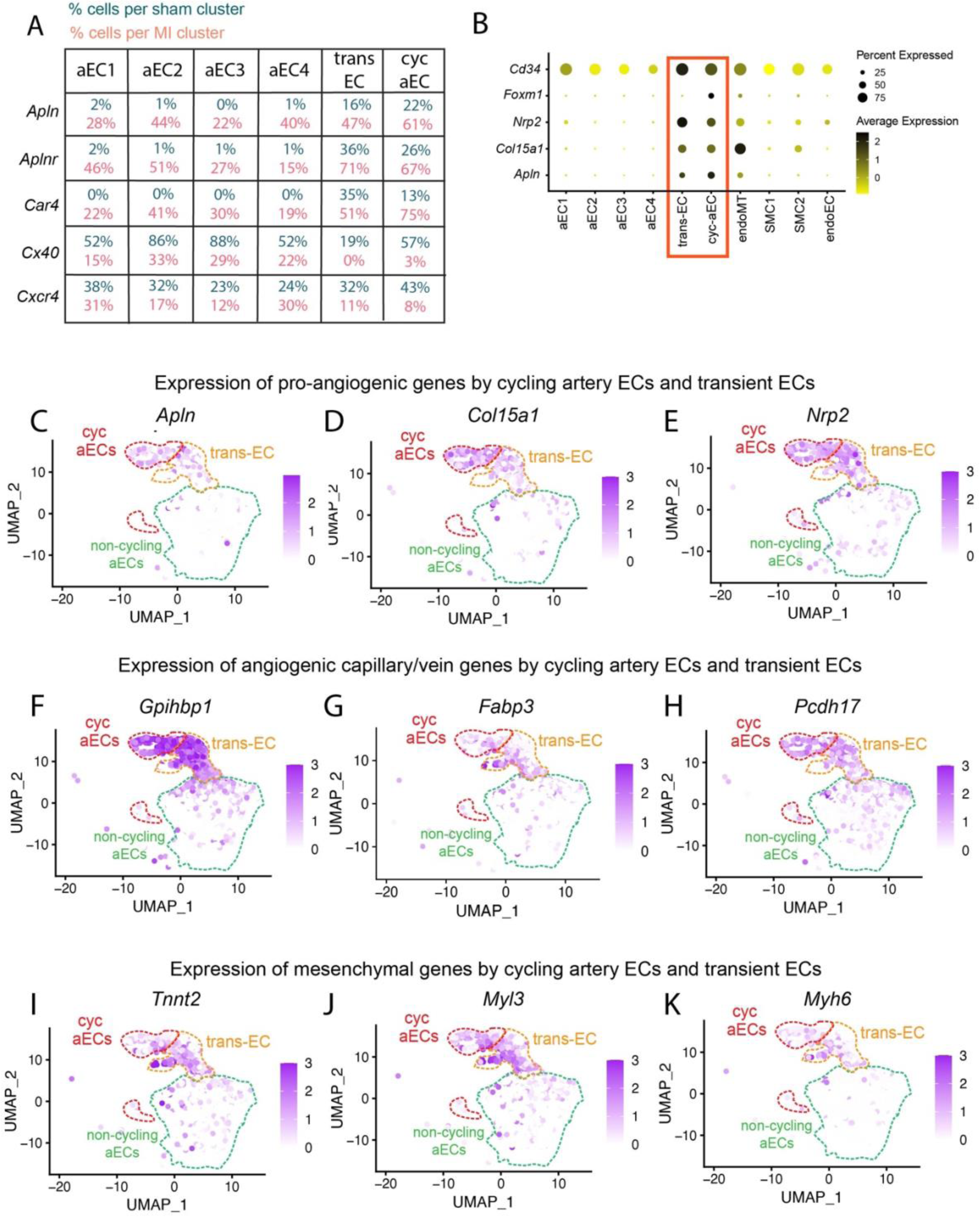
Analyses of de-differentiation features in *Cadherin5*^+^ neonatal cells. (**A**) Table showing percentage of sham and MI cells expressing *Apln*, *Aplnr*, *Car4*, *Cx40* and *Cxcr4*. (**B**) DotPlot from neonatal *Cadherin5*^+^ cells showing average expression and percentage of cells expressing pro-angiogenic genes (*Cd34*, *Foxm1*, *Nrp2*, *Col15a1*, *Apln*). (**C**-**E**) Feature Plots showing expression of pro-angiogenic *Apln*, *Col15a1*, and *Nrp2* in *Cadherin5*^+^ neonatal cells, from sham and MI together. (**F**-**H**) Feature Plots showing expression of angiogenic genes, also expressed by capillary or venous ECs (*Gpihbp1*, *Fabp3*, *Pcdh17*), in *Cadherin5*^+^ neonatal cells, from sham and MI together. (**I**-**K**) Feature Plots showing expression of mesenchymal genes (*Tnnt2*, *Myl3*, *Myh6*), in *Cadherin5*^+^ neonatal cells, from sham and MI together.

**Supplemental Figure 7:**
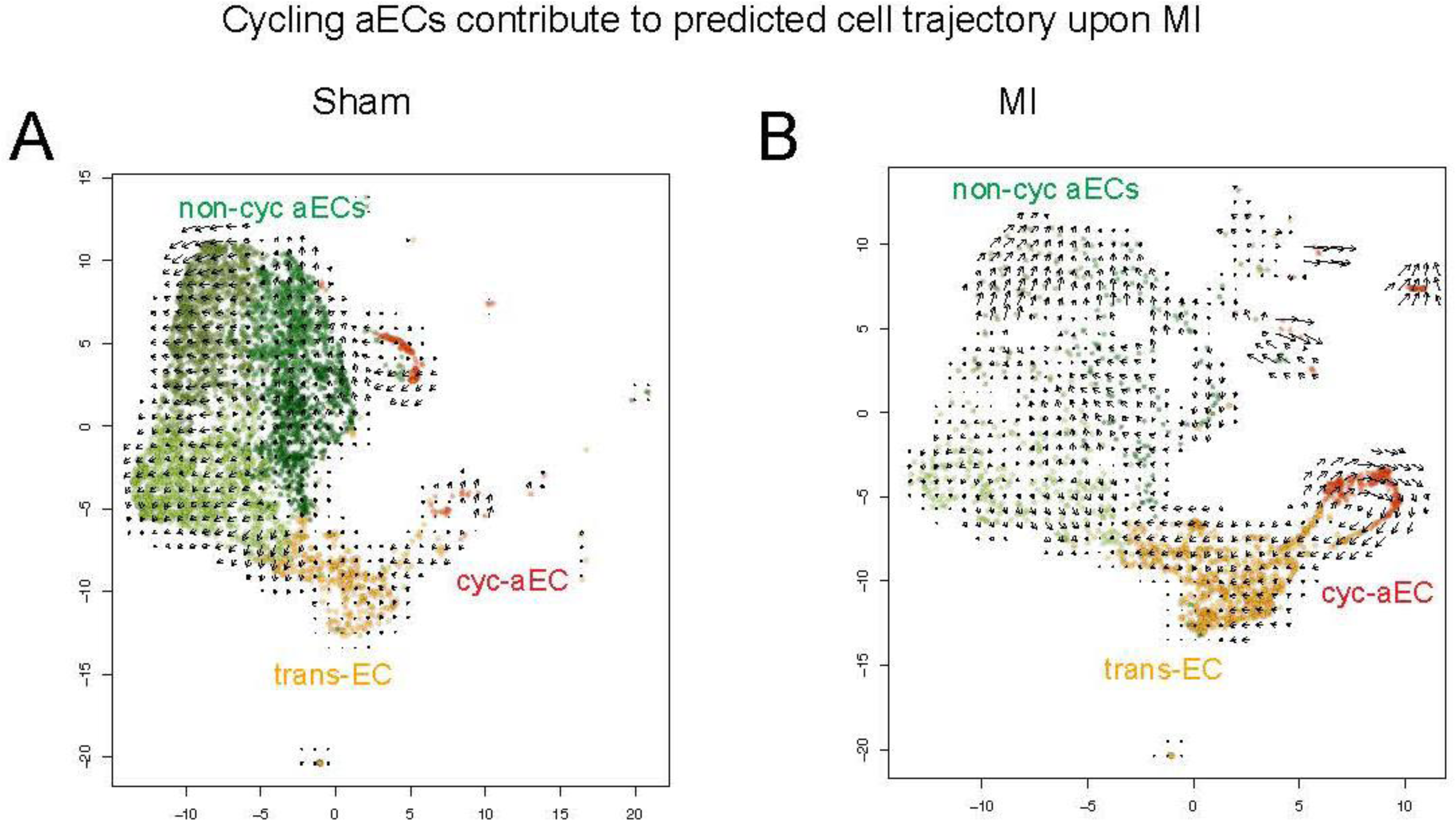
Comparison of de-differentiation features between neonatal *Cadherin5*^+^ cells in cyc-aEC, trans-EC and aEC1 clusters. (**A**, **B**) RNA velocity analysis on *Cadherin5^+^* (**A**) sham and (**B**) MI cells.

**Supplemental Figure 8:**
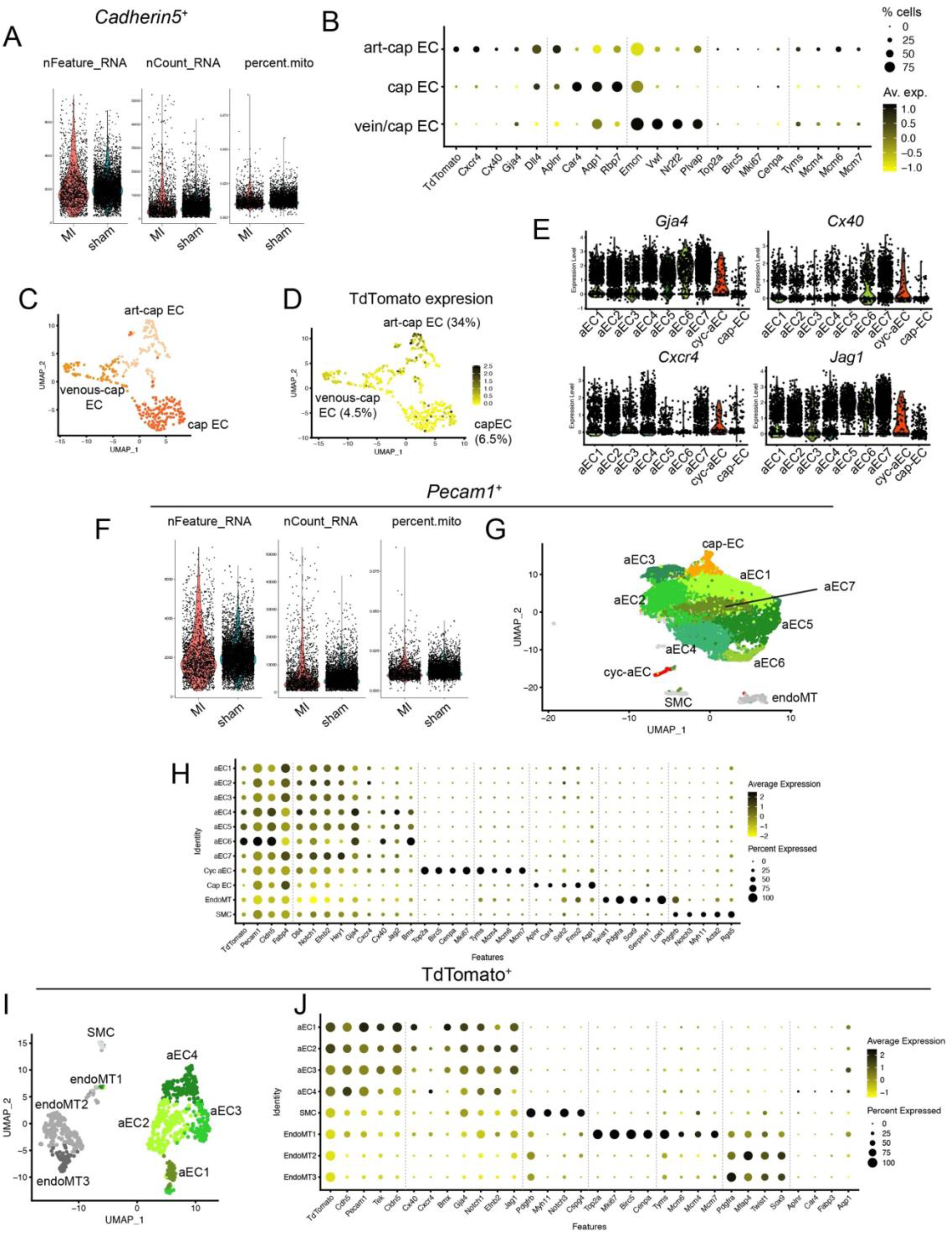
Identification of adult cardiac cell clusters. (**A**) Violin plots showing quality control (analyses of nFeature counts, nCount_RNA and mitochondrial genes) performed on adult *Cadherin5*^+^ dataset. (**B**) DotPlot showing percentage of cells and their average gene expression used for identifying clusters, obtained from re-clustering *Cadherin5*^+^ cap-ECs. (**C**) Visualization of re-clustered adult *Cadherin5*^+^ cap-ECs on a UMAP. Clusters were identified using the DotPlot shown in **B**. (**D**) FeaturePlot showing the distribution of TdTomato expressing cells on re-clustered cap-ECs in **C**. (**E**) Violin Plots showing expression of artery specific genes in adult *Cadherin5*^+^ cells in both sham and MI group, integrated. (**F**) Violin plots showing quality control (analyses of nFeature counts, nCount_RNA and mitochondrial genes) performed on adult *Pecam1*^+^ dataset. (**G**) Visualization of clusters in adult *Pecam1*^+^ cells, using UMAP. (**H**) DotPlot showing average gene expression and percentage of cells expressing cell type specific genes in *Pecam1*^+^ dataset. (**I**) Visualization of clusters in adult TdTomato^+^ cells, using UMAP. (**J**) DotPlot showing average gene expression and percentage of cells expressing cell type specific genes in TdTomato^+^ dataset.

**Supplemental Figure 9:**
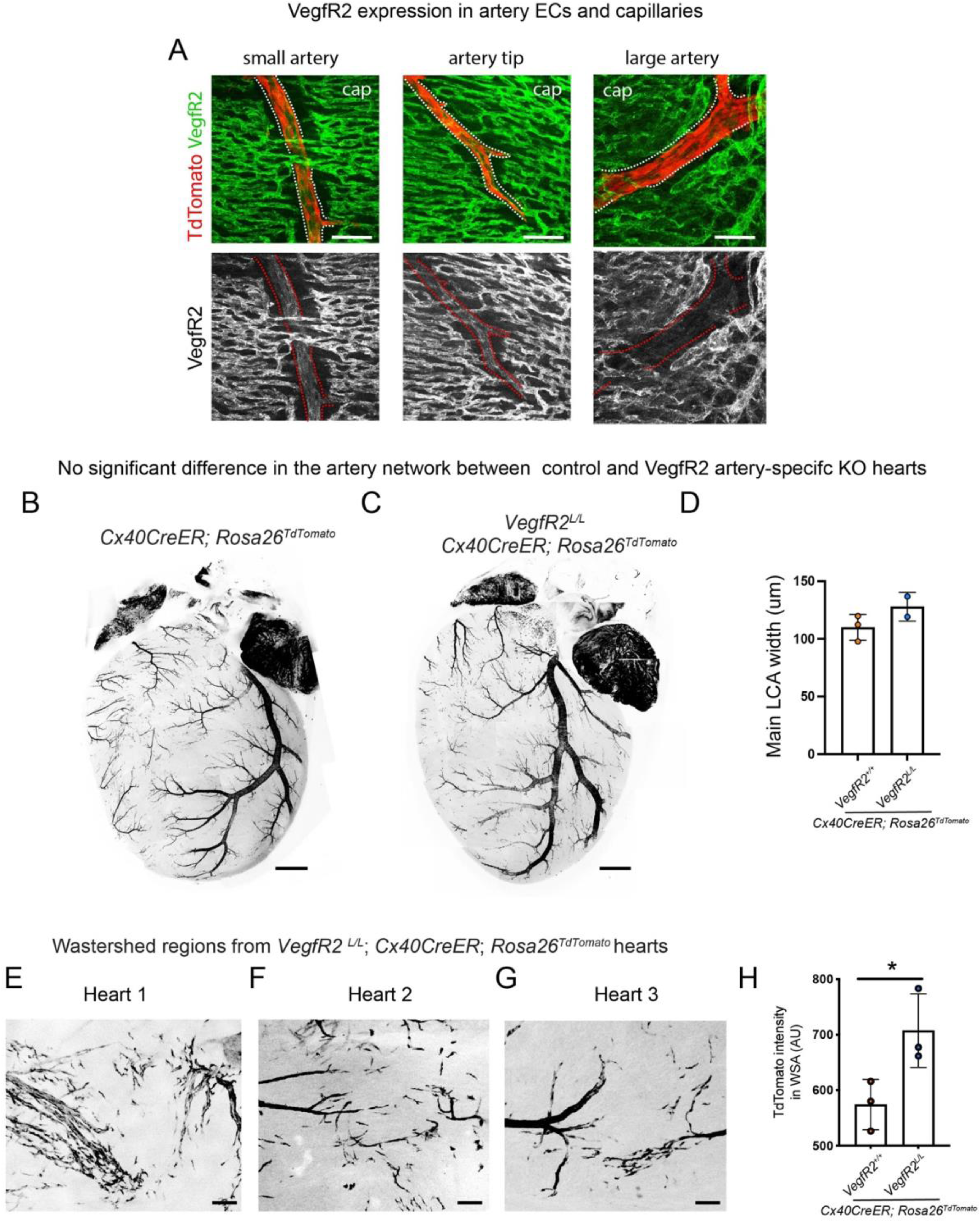
Analyses of arterial Vegf pathway in uninjured and injured neonatal hearts. (**A**) Representative confocal images from uninjured P6 *Cx40CreER*; *Rosa26^TdTomato^* neonatal hearts showing expression of VegfR2 in small arteries, artery tips and capillaries, but not in large arteries. (**B**-**D**) Representative whole heart images from uninjured *Cx40CreER*; *Rosa26^TdTomato^* mice, (**B**) with or (**C**) without arterial EC *VegfR2*. Tamoxifen was injected at P0, hearts were explanted at P6. (**D**) Quantification of width of main left coronary artery (LCA) in uninjured control and hearts deleted for *VegfR2* from arterial ECs. Tamoxifen was injected at P0, hearts were explanted at P6. The difference was statistically insignificant (p-value 0.1881). (**E**-**G**) Representative confocal images of *VegfR2* depleted watershed regions from P6 neonatal *Cx40CreER*; *Rosa26^TdTomato^* hearts, 4 days post-MI. Tamoxifen was administered at P0/P1, MI was performed at P2/P3. (**H**) Quantification of *Cx40CreER*-lineage traced (TdTomato^+^) saECs in *VegfR2* depleted watershed regions from P6 neonatal hearts (shown in **E**-**G**), 4 days post-MI. * represents p-value 0.0449. cap, capillary; KO, knockout; wsa, watershed area. Scale bars: **A**: 50µm; **B**, **C**: 500 µm; **E**-**G**: 100 µm

**Supplemental Figure 10:**
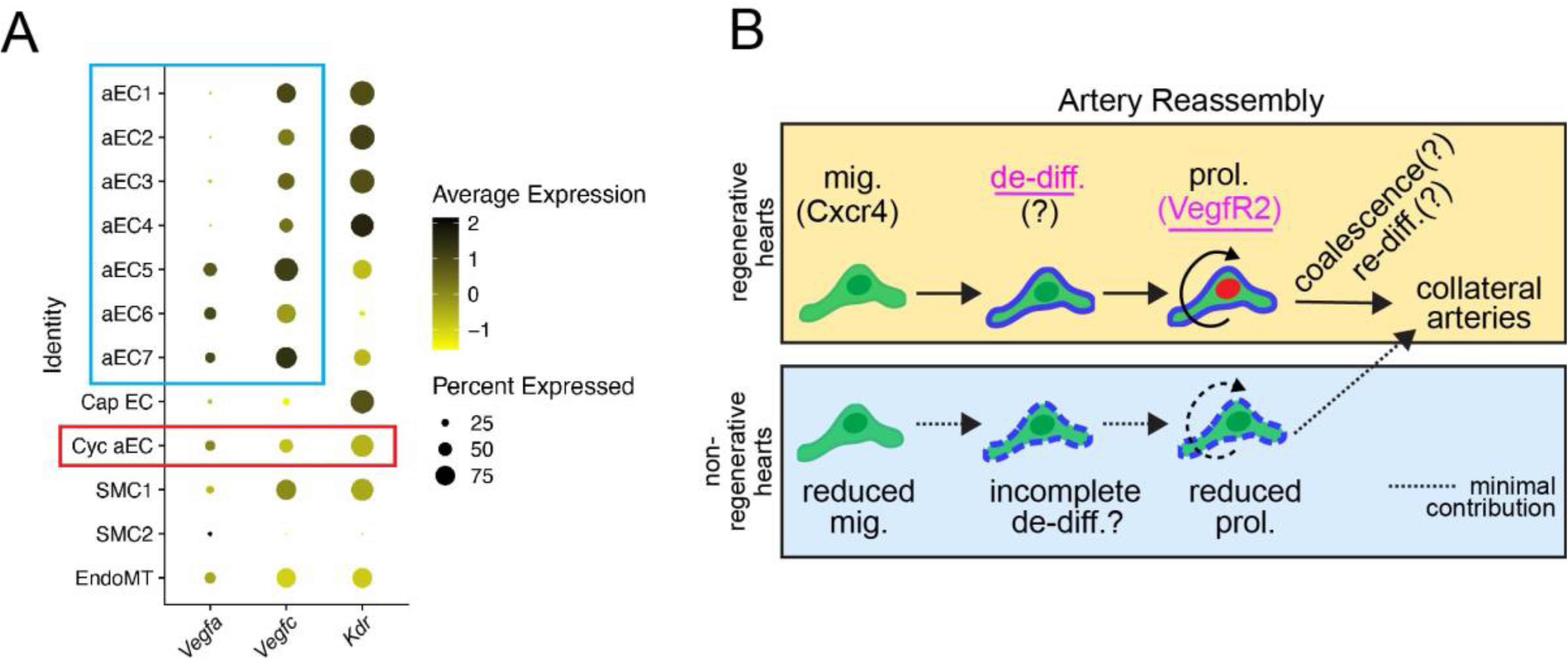
Proposed working model. (**A**) Dot plot showing average gene expression and percentage of cells expressing *Vegfa*, *Vegfc* and *VegfR2 (Kdr*) in adult *Cadherin5*^+^ dataset. Blue box indicates ligand expression by aECs, red box indicate downregulation of *VegfR2* in cyc-aEC cluster. (**B**) Proposed working model of cellular events associated with Artery Reassembly. In regenerative neonatal hearts, MI induces sequential migration (via Cxcr4), de-differentiation, and proliferation (via VegfR2) of pre-existing artery ECs, which, leads to formation of collateral arteries and efficient revascularization of ischemic heart regions. Non-regenerative hearts show minimal migration, de-differentiation or proliferation. While Cxcl12/Cxcr4 pathway drives migration, in this study we show that artery EC proliferation is regulated by Vegf pathway. Green, artery ECs; Blue, de-diff (de-differentiation); Red, prol. (proliferation); mig, migration;? unknown molecules associated with the cellular events linked to Artery Reassembly. Findings of this study (de-differentiation and VegfR2-regulated proliferation of artery ECs) are underlined.

**Table S1:** Summary of all p-values obtained from Wilcoxon sum rank test.

| Statistical significance of gene expression variation as observed using<br>Wilcoxon test |  |  |  |  |  |  |
| --- | --- | --- | --- | --- | --- | --- |
| Statistical analysis of de-differentiation genes across neonatal *Cadherin5*<sup>+</sup><br>sham and MI cyc-aEC clusters |  |  |  |  |  |  |
|  | p_val | auc | avg_log2FC | pct.1 | pct.2 | p_val_adj |
| *Gja5* | 6.80E-52 | 0.225561 | -1.39143 | 0.026 | 0.569 | 1.30E-47 |
| *Cxcr4* | 1.71E-20 | 0.329778 | -0.30521 | 0.082 | 0.431 | 3.27E-16 |
| *Aplnr* | 1.75E-17 | 0.727798 | 0.699507 | 0.673 | 0.257 | 3.36E-13 |
| *Car4* | 5.90E-06 | 0.598174 | 0.413379 | 0.312 | 0.125 | 1.13E-01 |
| *Apln* | 1.57E-14 | 0.701489 | 0.762624 | 0.608 | 0.222 | 3.02E-10 |
| *Foxm1* | No significant difference |  |  |  |  |  |
| Statistical analysis of de-differentiation genes across neonatal *Cadherin5*<sup>+</sup> MI<br>cyc-aEC and trans-EC clusters |  |  |  |  |  |  |
|  | p_val | auc | avg_log2FC | pct.1 | pct.2 | p_val_adj |
| *Gja5* | No significant difference |  |  |  |  |  |
| *Cxcr4* | No significant difference |  |  |  |  |  |
| *Aplnr* | 4.82E-12 | 0.377149 | -0.5043 | 0.673 | 0.758 | 9.23E-08 |
| *Car4* | 1.02E-07 | 0.415964 | -0.55117 | 0.312 | 0.436 | 1.95E-03 |
| *Apln* | No significant difference |  |  |  |  |  |
| *Foxm1* | 7.04E-44 | 0.654272 | 0.367115 | 0.341 | 0.031 | 1.35E-39 |
| **Statistical analysis of de-differentiation genes across neonatal *Cadherin5*<sup>+</sup> MI** |  |  |  |  |  |  |
| **trans-EC and aEC1 clusters** |  |  |  |  |  |  |
|  | p_val | auc | avg_log2FC | pct.1 | pct.2 | p_val_adj |
| *Gja5* | 1.64E-24 | 0.423338 | -0.30514 | 0.001 | 0.155 | 3.14E-20 |
| *Cxcr4* | 5.20E-12 | 0.398321 | -0.40858 | 0.117 | 0.314 | 9.97E-08 |
| *Aplnr* | 2.26E-16 | 0.684601 | 0.517376 | 0.758 | 0.367 | 4.33E-12 |
| *Car4* | 1.43E-08 | 0.613955 | 0.616043 | 0.436 | 0.222 | 2.74E-04 |
| *Apln* | 7.20E-22 | 0.694867 | 0.94641 | 0.488 | 0.106 | 1.38E-17 |
| *Foxm1* | No significant difference |  |  |  |  |  |
| **Statistical analysis of de-differentiation genes across neonatal *Cadherin5*<sup>+</sup> MI** |  |  |  |  |  |  |
| **cyc-aEC and aEC1 clusters** |  |  |  |  |  |  |
|  | p_val | auc | avg_log2FC | pct.1 | pct.2 | p_val_adj |
| *Gja5* | No significant difference |  |  |  |  |  |
| *Cxcr4* | 1.00E-14 | 0.378972 | -0.48813 | 0.082 | 0.314 | 1.92E-10 |
| *Aplnr* | No significant difference |  |  |  |  |  |
| *Car4* | No significant difference |  |  |  |  |  |
| *Apln* | 5.76E-30 | 0.753943 | 1.028363 | 0.608 | 0.106 | 1.10E-25 |
| *Foxm1* | 2.78E-19 | 0.663357 | 0.399056 | 0.341 | 0.014 | 5.33E-15 |
| **Statistical analysis of de-differentiation genes across adult *Cadherin5*<sup>+</sup> sham and MI cyc-aEC clusters** |  |  |  |  |  |  |
|  | p_val | auc | avg_log2FC | pct.1 | pct.2 | p_val_adj |
| *Gja5* | No significant difference |  |  |  |  |  |
| *Cxcr4* | 0.037701 | 0.373907 | -0.57171 | 0.327 | 0.5 | 1 |
| *Aplnr* | 0.015657 | 0.612245 | 0.824505 | 0.265 | 0.036 | 1 |
| *Car4* | No significant difference |  |  |  |  |  |
| *Apln* | 0.091189 | 0.565233 | 0.623217 | 0.163 | 0.036 | 1 |
| *Foxm1* | No significant difference |  |  |  |  |  |
| **Statistical analysis of *Vegf* pathway genes across neonatal *Cadherin5*<sup>+</sup> sham and MI cyc-aEC clusters** |  |  |  |  |  |  |
|  | p_val | auc | avg_log2FC | pct.1 | pct.2 | p_val_adj |
| *Kdr* | 3.49E-22 | 0.770358 | 0.799244 | 0.933 | 0.743 | 6.69E-18 |
| *Vegfa* | 1.53E-21 | 0.299646 | -0.49758 | 0.144 | 0.528 | 2.93E-17 |
| *Vegfc* | 1.81E-51 | 0.176933 | -1.4309 | 0.099 | 0.708 | 3.48E-47 |
| **Statistical analysis of *Vegf* pathway genes across adult *Cadherin5*<sup>+</sup> sham and MI cyc-aEC clusters** |  |  |  |  |  |  |
|  | p_val | auc | avg_log2FC | pct.1 | pct.2 | p_val_adj |
| *Kdr* | 1.45E-15 | 0.125061 | -1.96031 | 0.857 | 0.97 | 2.67E-11 |
| *Vegfa* | 3.01E-03 | 0.586006 | 0.375123 | 0.286 | 0.107 | 1.00E+00 |
| *Vegfc* | 1.29E-02 | 0.586856 | 0.265401 | 0.388 | 0.19 | 1.00E+00 |

